# Uncertainty underlies geometric effects on navigation and grid-cell tuning

**DOI:** 10.1101/2023.01.30.526278

**Authors:** Yul HR Kang, Daniel M Wolpert, Máté Lengyel

## Abstract

Knowing one’s place in the world requires integrating sensory inputs with respect to the geometry of the environment. Consistent with this, variations in environmental geometry, such as the shape and size of an enclosure, have profound effects on navigational behavior^1–4^ and its neural underpinning in grid cells^5–7^. Here, we show that these effects arise as a consequence of a single, unifying principle: to navigate efficiently, the brain must maintain and update the uncertainty about one’s location. We develop an image-computable Bayesian ideal observer model of navigation, continually combining noisy visual and self-motion inputs, and a neural encoding model representing the spatial uncertainty computed by the ideal observer. Critically, we find that a key determinant of spatial uncertainty is the dimensionality reduction inherent in the retinal projection of the environment. Mathematical analysis and simulations show that spatial uncertainty accounts for a diverse range of sometimes paradoxical distortions in human homing behavior across trapezoidal, stretched, and compressed environments. Moreover, the neural encoding of this uncertainty accounts for observed changes in grid field size, anisotropy, rescaling, and boundary-dependent tethering under analogous geometric manipulations. Our results show that spatial uncertainty arising unavoidably during navigation is key to understanding navigational behavior and its neural underpinnings.

---

Determining our location in the environment is essential for survival^8^. This process engages well-known neural populations, including hippocampal place cells and entorhinal grid cells^9,10^, the latter forming periodic firing patterns proposed to provide a spatial metric^11^. Yet, these responses – and navigational behavior – are distorted when the environment’s geometry is altered. For example, in trapezoidal enclosures, human spatial memory becomes biased^1,12^, and grid cell tuning becomes less regular^5,12^. When environments stretch or compress along one axis, both behavior and grid responses show unexpected distortions even along the unchanged axis^2,3,6^. These effects can also depend on the most recently approached wall, a phenomenon known as tethering^4,7^. While these distortions may sometimes appear small and idiosyncratic, here we propose that they reveal a critical and unifying principle of navigation: that the brain continuously represents its uncertainty about spatial location as fundamentally uncertain, and this uncertainty drives both behavior and neural responses. Unlike previously used simplified landmark-based inputs^13–17^, we use an image-computable model to determine the spatial uncertainty arising from the two-dimensional retinal projection of a threedimensional environment and its optimal integration with self-motion inputs. We show that this spatial uncertainty accounts for key effects in how environmental geometry shapes behavior and neural responses that were previously explained by distinct and often heuristic processes^1–4,7,12,18–22^.

## The BION model

To study spatial uncertainty, we formalized the problem of location estimation as recursive probabilistic inference over allocentric pose (location and heading) based on egocentric sensory signals – visual input (modeled as a retinal image; Supp. Material SM1), self-motion cues such as vestibular input (translation and rotation) and, for simulating rat navigation, tactile input from nearby walls – using an internal model of how these variables are related during navigation (Fig. 1a-b, Methods).

**Fig. 1.**
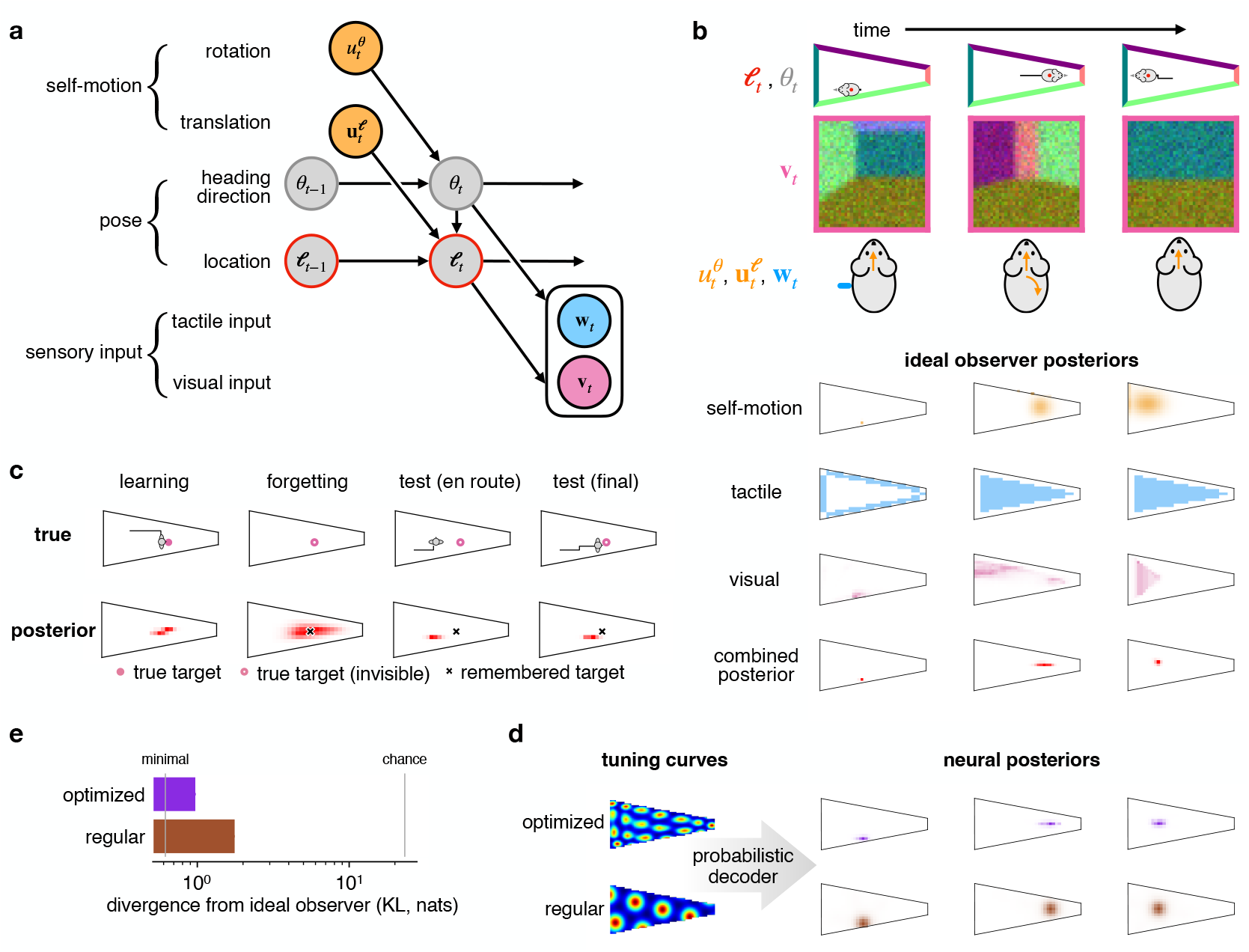
The BION model. **a**. Generative model: allocentric location (ℓ*t*) and heading (*θ*_*t*_) are updated via self-motion inputs 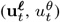, yielding visual (**v**_*t*_) and tactile (**w**_*t*_) observations. Gray/colored circles: hidden/observed variables. **b**. *Row 1:* Simulated trajectory of a rat (1^st^ –3^rd^ columns). *Row 2:* Visual input (**v**_*t*_) with blur and noise. *Row 3:* Self-motion (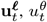, orange) and tactile (**w**_*t*_, blue: location of touch on body) inputs. *Rows 4–7:* Ideal observer posteriors over location (ℓ_*t*_): *row 4*, self-motion only, 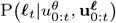; *row 5*, tactile only, P(**w**_*t*_|ℓ_*t*_); *row 6*, visual only, P(**v**_*t*_|ℓ_*t*_); *row 7*, all cues combined,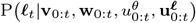. **c**. Behavioral model: memory encodes noisy posterior mean (black×) after learning (1^st^–2^nd^ columns); test phase simulates movement toward remembered location by minimizing expected squared error under current posterior (3^rd^–4^th^ columns). **d**. Neural model: responses from regular (bottom) or warped (top) grid fields (left) decoded into neural posteriors (right), matched to ideal observer posteriors (**b**, row 7) via KL divergence minimization. **e**. Mean KL divergence over time: regular (brown) and optimized (purple). Vertical lines show controls for the minimal achievable and the chance-level (time-shuffled) KL (Methods). Error bars: s.e.m.

We used this Bayesian Image-computable Observer for Navigation (BION) model to predict behavioral responses in human spatial memory tasks and grid cell responses in rodents (Methods). We focused on paradigms where both behavioral and neural data were available (Extended Data Figs. 1 and 2), and where participants were well-familiarized with the environment to minimize uncertainty about the layout^16^, isolating location uncertainty within a known map. In the human experiments, participants performed homing tasks in virtual reality, returning to previously viewed objects’ locations. We modeled both the learning and test phases of these experiments, including forgetting between them (Fig. 1c).

To model grid cell responses, we simulated a rat foraging and generated neural activity from a population of grid cells. A probabilistic population decoder^23,24^ was used to recover the neural posterior over location (Fig. 1d, right; Methods). Instead of minimizing uncertainty in the neural posterior as in standard efficient coding models^25^, we required it to match the uncertainty of the ideal observer. In particular, the ideal observer’s posterior changed dynamically over time, with ongoing modulations of the overall extent and anisotropy of its uncertainty (Fig. 1b, row 7). Grid cells with standard hexagonal tuning showed translation-invariant, isotropic neural posteriors that failed to capture these modulations (Fig. 1d, bottom; discrepancy quantified in Fig. 1e, brown). To match the ideal observer, we implemented a ‘warped population code’^21,25^, adjusting the grid structure so that decoded neural posteriors aligned with the ideal observer’s moment-by-moment uncertainty (Fig. 1d, top; Fig. 1e, purple; Extended Data Fig. 3), resulting in neural responses (approximately) encoding the ideal observer’s posteriors in an easily decodable format (Supp. Material SM2, Extended Data Fig. 4). These warped responses provided our predictions for grid cell tuning. Thus, rather than explaining the origin of hexagonal symmetry as in prior models^26–32^, we show how symmetry degrades under environmental manipulations to improve the encoding of uncertainty.

### Deformations in familiar environments arise from inhomogeneous uncertainty

A critical analytical insight about how images are projected onto the retina is that the same landmark (e.g. a corner of the enclosure) provides different amounts of visual information about the participant’s distance from it depending on its distance along, *d*, and eccentricity from, *ϵ*, the participant’s line of sight (Fig. 2a, inset; Supp. Material SM3):

**Fig. 2.**
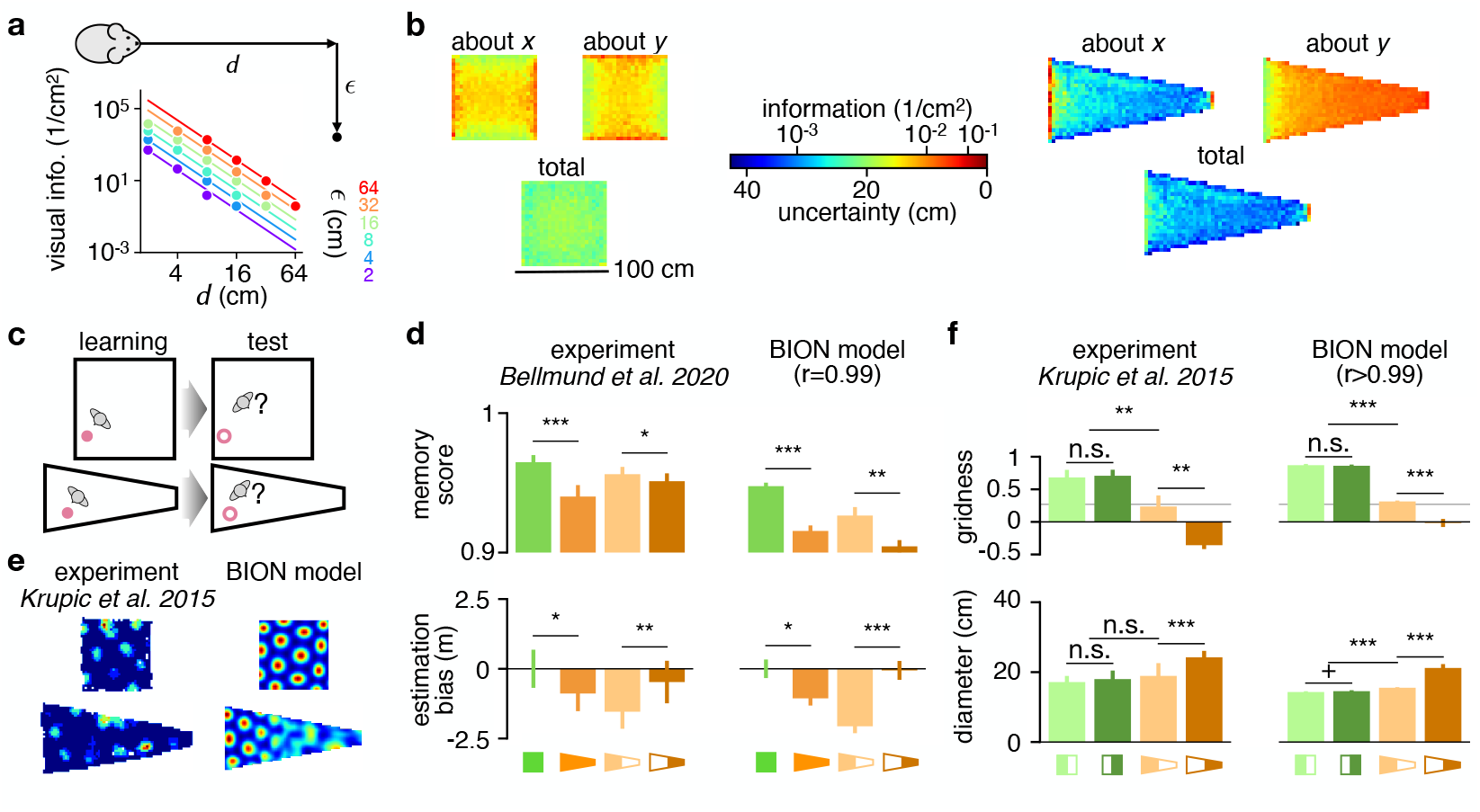
Deformations in familiar environments arise from inhomogeneous uncertainty. **a**. Inset: top-down view of an observer and a landmark, defined by distance along (*d*) and eccentricity from line of sight (*ϵ*). Main: Fisher information about *d* vs. true *d* and *ϵ* (log–log scale); lines: analytic (Eq. 1), markers: simulation. **b**. Visual information about *x, y*, and total location information in square vs. trapezoid environments (hotter = more info). **c**. Homing task^1^ in square (top) and trapezoid (bottom; cf. Fig. 1c). **d**. Spatial memory in data^1^ (left) and model (right). Columns: square vs. trapezoid; broad vs. narrow halves. Top: memory score (target rank). Bottom: distance estimation bias. Error bars: s.e.m. across participants. **e**. Example grid fields in data^5^ (left) and BION (right) in square (top) vs. trapezoid (bottom). **f**. Gridness (top) and field diameter (bottom) by environment half in data^5^ (left) and BION (right). Gray line indicates the inclusion threshold in the square environment. Error bars: s.e.m. across cells. Statistical tests follow original experiments. Significance: n.s. *p>*0.10, + *p<*0.10, * *p<*0.05, ** *p<*0.01, *** *p<*0.001; r-values: model–experiment correlations (allowing offset/scaling).

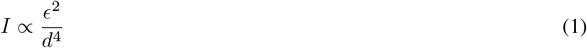

This equation captures the intuition that, the closer and more eccentric a landmark is, the more its retinal image shifts per unit movement, providing information about distance. Indeed, numerical simulations of the full image-computable BION model confirmed these calculations (Fig. 2a).

In line with Eq. 1, BION makes direct predictions about how visual information about location changes inside a squareversus a trapezoid-shaped environment, such as those used in several experiments^1,5,12^ (Fig. 2b, Extended Data Fig. 5a). In turn, the inhomogeneity of these visual information maps explains a range of experimental results. In human homing tasks (Fig. 2c, cf. Fig. 1c), total errors are dominated by the more uncertain direction, i.e. position about the long axis in the trapezoid (Fig. 2b, compare top two panels for the trapezoid, showing uncertainty about x and y separately, with the bottom panel, showing total uncertainty). Along this direction, on average, *d* is greater and thus information is smaller than along either direction in the square. Therefore, BION predicts homing performance to be worse in the trapezoid than in the square, in line with experimental findings^1^ (Fig. 2d, top, green vs. orange). Finally, the asymmetry between the two ends of the trapezoid results in *ϵ* being larger and thus information being greater near the broad end. As a consequence, homing performance is predicted to be relatively better near its broad end than near the narrow end, again consistent with data^1^ (Fig. 2d, top, light vs. dark brown).

Participants also estimated distances between previously visited objects. Their estimation biases did not simply reflect their memory performance for individual locations: while the biases in the trapezoid were overall more negative than in the square (in line with memory performance being poorer in the trapezoid than in the square), they were *less* negative (although memory performance was *more* diminished) in the narrow side of the trapezoid than in the broad side (Fig. 2d, left, compare bottom to top). BION reproduced this pattern of results (Fig. 2d, bottom right). Our analysis revealed that this effect is also explained mathematically by the principle of Bayesian cue combination when information provided by visual cues is spatially inhomogeneous as suggested by Eq. 1 (Supp. Material SM4).

Because grid cell responses in BION encode spatial uncertainty, the size and anisotropy of predicted grid fields respectively depended on the overall level and anisotropy of the ideal observer’s spatial uncertainty (Fig. 1d, and Fig. 2e, right; Extended Data Fig. 6). Compared to the square enclosure, uncertainty in the trapezoid was greater and more anisotropic (differing greatly along the two axes of the enclosure, Fig. 2b). Thus, in line with experimental measurements^5^ (Fig. 2e–f, left), BION predicted larger grid fields with lower levels of gridness for the trapezoid than for the square (Fig. 2e, right and f, right, brown bars vs. green bars), and for the narrow than for the broad side of the trapezoid (Fig. 2f, right, dark vs. light brown), and generally lower similarity of grid fields across the two sides of the environment in the trapezoid than in the square enclosure (Extended Data Fig. 5b).

Overall, BION accurately accounted for the qualitative effects of a trapezoid vs. a square enclosure both on human behavior (r=0.99, Methods) and on grid cell responses (r>0.99).

### Scaling and biases in novel environments arise from model mismatch

Next, we asked if BION could also explain the scaling of behavioral response patterns when rectangular environments were expanded or compressed along one or both axes (Fig. 3a–b, Extended Data Fig. 7). Because the change of the environment in the test phase was unknown to participants, we modeled the ideal observer’s inferences as being computed in the familiar environment’s frame of reference, resulting in a model mismatch between the internal model used by the ideal observer and the actual environment (see Methods). In line with experimental data^2^, this resulted in higher uncertainty and thus more response variability along the axis of the environment that was changed (Fig. 3b, axes with arrowheads), and for locations along the axis for which information was low to begin with (because they were farther from the nearest wall; Fig. 3b, red vs. blue arrowheads).

**Fig. 3.**
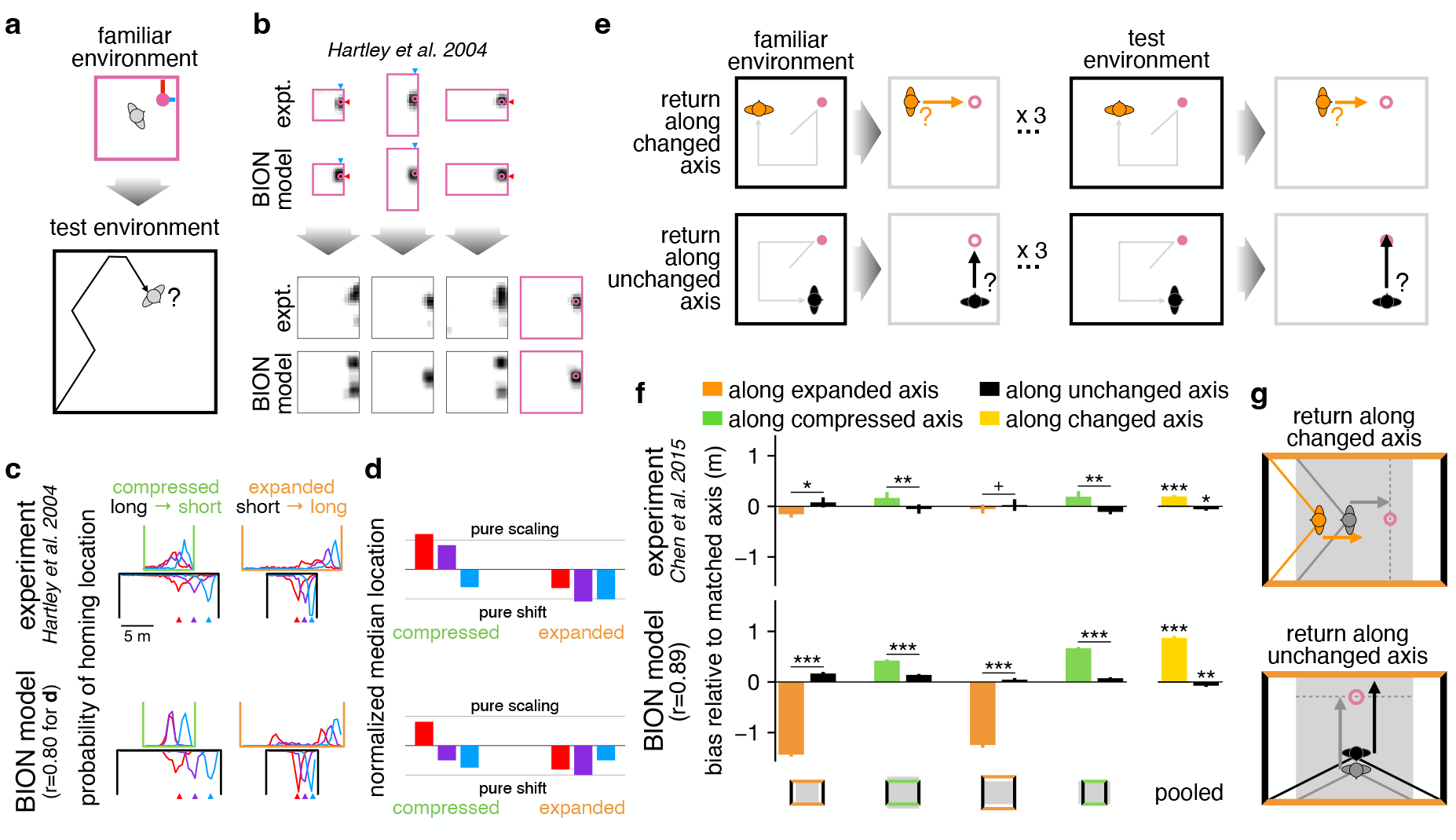
Scaling and biases of human behavior in novel environments arise from model mismatch. **a**. Homing in scaled environments^2^. Pink circle: learned target; blue/red lines: distances to nearest walls along horizontal/vertical axes. **b**. Homing location distributions in familiar (pink-outlined) and test (black-outlined) environments in data^2^ (rows 1,3) and BION (rows 2,4). Blue/red arrowheads mark target distances that are shorter/longer from nearest wall (cf. **a**). See Extended Data Fig. 7 for all conditions. **c**. Distributions along scaled (upright) vs. unchanged (mirrored) axes for compressed (left) and expanded (right) test environments, for near (blue), intermediate (purple), and far (red) target-wall distances in data^2^ (top) and BION (bottom). **d**. Modes of distributions from **c**, normalized between pure scaling (top line) and pure wall-based shifting (bottom line) in data^2^ (top) and BION (bottom). **e**. Axis-dependent homing^3^. Pink circle: target; gray line: outbound paths; gray bounding boxes/pink unfilled circles: no visual cues; orange/black arrows: return along changed/unchanged axis. **f**. Homing biases (±s.e.m. across participants) along changed (orange/green) and unchanged (black) axes in data^3^ (top) and BION (bottom) corresponding to the conditions shown in **e** (Experiments 1 & 2 of Ref. 3). Schematics show familiar (gray) and test (black/colored) environments. Yellow/black bars: pooled average (± s.e.m. across experiments) with changed/unchanged axes. Positive: undershoot in compression or overshoot in expansion. See full results in Extended Data Fig. 8. **g**. Schematic of how biases arise when the test environment is scaled (orange: changed, black: unchanged axis) compared to the familiar environment (gray). Significance levels in **f** and r-values in **c**–**d** and **f** are shown as in Fig. 2.

There were also systematic biases in homing responses in the same experiment. In most cases, the distributions of actual homing locations were better explained by shifting with the closest wall than by scaling with the environment, except for targets farther from the walls when the environment was compressed (Fig. 3c–d, top). BION reproduced these results with good accuracy (Fig. 3c–d, bottom, r=0.80). These effects, along with the occasional bimodal homing response distributions seen in expanded environments (Fig. 3b), can also be derived from the location-dependent asymmetric informativeness of visual cues supplied by different walls, generally making visual cues from nearer walls dominate and thus responses to shift with them (Eq. 1, Supp. Material SM5).

The same principle of model mismatch also accounted for conflicting response biases along the changed and unchanged axes of a scaled environment^3^ (Fig. 3e-f; Extended Data Fig. 8). First, responses were negatively biased (i.e. undershot) along an expanded axis (Fig. 3f, top, orange bars) and positively biased (i.e. overshot) along a compressed axis (Fig. 3f, top, green bars). Second, responses along the unchanged axes were biased, and in the opposite direction: positively (overshooting) when the environment was expanded, and negatively (undershooting) when it was compressed (Fig. 3f, top, black bars). BION captured both these kinds of biases. First, just before homing starts, the visual image of the far wall along the expanded axis suggests that this wall is still at a great distance, which in turn makes the participant’s distance to the target seem shorter than it actually is, leading to an undershoot in simulated responses (Fig. 3f, bottom, orange and Fig. 3g, top) – and vice versa when the far wall is seen along the compressed axis (Fig. 3f, bottom, green). Second, looking along the unchanged direction, the farther expanded wall subtends a large angle making it appear as if it was closer, leading to an overshoot in simulated responses (Fig. 3g, bottom) – and again vice versa in the compressed environment (Fig. 3f, bottom, black bars). Therefore, when pooled across conditions, the sign of the response bias along the changed vs. unchanged axis was respectively the opposite vs. the same as the sign of the change in the environment’s size (Fig. 3f, ‘pooled’), and BION provided a good overall match to experimental data (Fig. 3f, Extended Data Fig. 8, r=0.89).

The anisotropy of grid cell responses in analogous experiments was found to depend on whether the shape of the enclosure was familiar or encountered after scaling, rather than on the shape itself (Fig. 4a, b top). In BION, grid cell responses still encode neural posteriors in the familiar enclosure (consistent with the ideal observer), while they are being recorded in the novel, scaled enclosure. Thus, the ideal observer’s distortions of uncertainty due to model mismatch that explained human behavior also resulted in increased warping of grid fields after environmental scaling, in line with experimental observations (Fig. 4a–b, bottom; Extended Data Fig. 9).

**Fig. 4.**
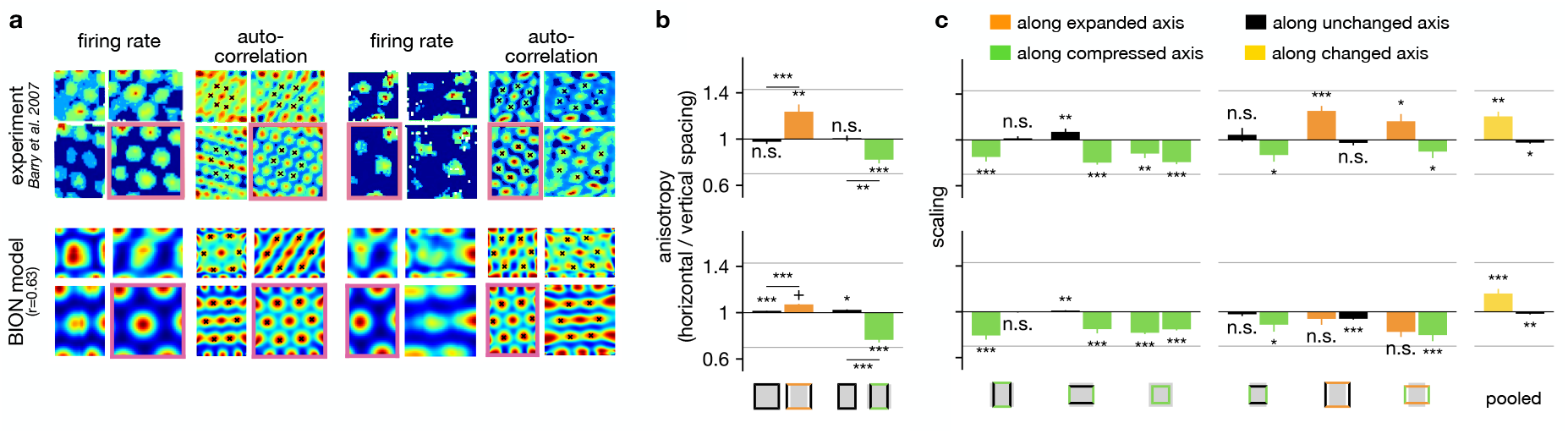
Scaling of rat grid fields in novel environments arises from model mismatch. **a**. Effect of environmental scaling on grid cells in the data^6^ (top) and BION (bottom). Rats explored a familiar room (pink outline; square vs. rectangular, columns 1–2 vs. 3–4) before scaled versions (three adjacent enclosures). For each experiment, the firing rate (columns 1,3) and autocorrelation map (columns 2,4) of a cell is shown. **b**. Grid anisotropy in data^6^ (top) and BION (bottom). Schematics show familiar (gray) and test (colored) boundaries; black, green, and orange indicate unchanged, compressed, and expanded axis. **c**. Grid scaling along changed (green/orange) and unchanged (black) axes in data^6^ (top) and BION (bottom). Rightmost bars pool results across all scaled environments, relative to their actual scaling (Methods). Gray horizontal lines indicate the anisotropy (**b**) and scaling (**c**) expected when the grid fields scale proportionally with the environment. Error bars: s.e.m. across cells. Significance and r-values as in Fig. 2.

Consistent with the biases seen in human behavior (Fig. 3f, top), rat grid fields compressed and expanded respectively along the compressed and expanded axes of the environment (Fig. 4c, top, green and orange bars), and showed opposite scaling along the unchanged direction (Fig. 4c, top, black bars). The same distortions in the BION model’s spatial uncertainty that explained human behavior also resulted in grid field warpings that reproduced these neural response patterns (Fig. 4c, bottom; r=0.63 for results shown in Fig. 4b–c).

### Tethering arises from biased cue combination

Further analyses of human homing responses in scaled environments revealed an intriguing ‘tethering’ effect with respect to the wall from which participants started their return journeys^4^ (Fig. 5a). While homing locations tended to compress or expand with the environment (Fig. 5b, left), as expected, they also showed systematic shifts (Fig. 5b, right): they shifted forwards or backwards, when the environment was compressed or expanded, respectively. Thus, homing locations tended to remain at a fixed distance from the starting wall (i.e. as if they were ‘tethered’ to it)^4^.

**Fig. 5.**
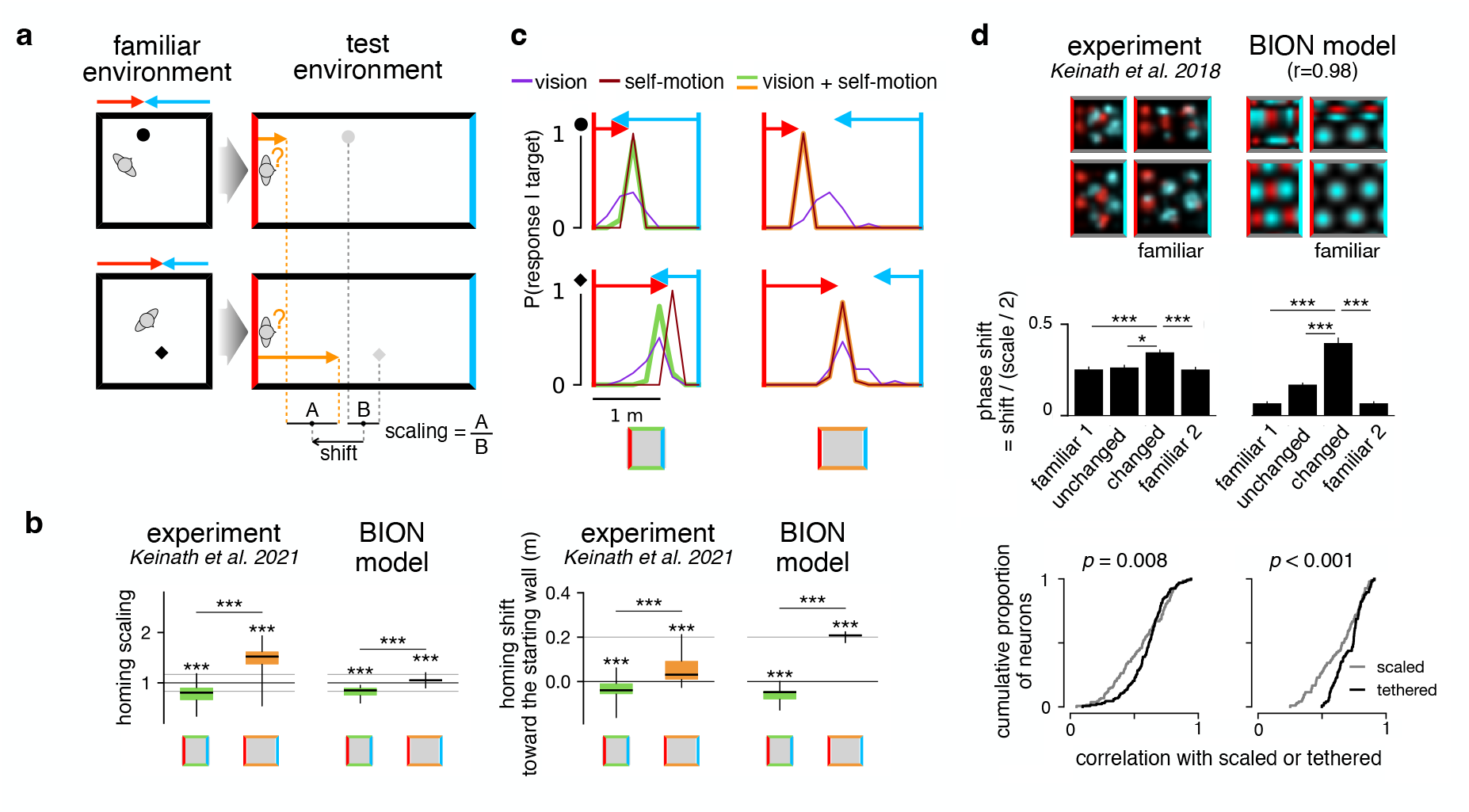
Tethering arises from biased cue combination. **a**. Distance-dependent homing in scaled environments^4^ (aspect ratios exaggerated). Left, black circle/diamond: learned target locations in square familiar environment near/far the starting wall used during test; red/blue arrows: target distances from starting/opposite wall used during test. Right, gray shapes: equivalent target locations in scaled test environment (same distance from center as in familiar environment); red/blue: starting/opposite wall of return journey to remembered target location. **b**. Scaling and shift in the data^4^ (left in each pair) and BION (right), shown as box-and-whisker plots. Gray lines show what would be expected from perfect scaling (left) or perfect tethering (right). **c**. Homing location distributions in BION for compressed (left) and expanded (right) environments, for targets near (top) or far (bottom) the starting wall based on vision only (purple), self-motion only (dark red), and both vision and self-motion (green, orange). Red/blue lines and arrow: starting/opposite walls and target distances from these walls in the familiar environment. Schematics show familiar (gray) and test (colored) boundaries. **d**. Grid fields in data^7^ (left) and BION (right). Top: Example firing rate maps conditioned on last approached wall (red: west; cyan: east) in four environments. Middle: Grid phase shifts relative to grid scale in familiar and compressed environments along unchanged and changed axis. Results are pooled across all four last approached walls (see Extended Data Fig. 11). Bottom: Cumulative distributions of Pearson’s correlations between observed tuning and those predicted by perfect scaling (gray) or shifting (black); p-values from Kolmogorov-Smirnov test. Significance and r-values in **d** middle as in Fig. 2.

BION explained these results as a fundamental feature of the ongoing multisensory integration of self-motion and visual inputs with asymmetric informativeness – arising from their respective route- and location-dependence. Self-motion inputs anchor the estimated location to the starting wall with high precision as the subject moves away from there, following a short, straight path, as in the experiments (Fig. 5c, brown distributions). Visual cues of the starting and opposite walls are in conflict, as they anchor the estimated location to the corresponding wall. Thus, depending on heading direction en route, one or the other set of visual cues, or their combination will determine the final homing location – resulting in a broader distribution of homing locations (Fig. 5c, purple distributions). Overall, ongoing integration of self-motion and visual inputs anchored to either wall results in interpolation between locations anchored to these walls, explaining the scaling component in homing responses (Fig. 5b, left). However, this interpolation is heavily biased: both self-motion input and visual cues from the starting wall support tethering to that wall. In addition, uncertainty from path integration only grows modestly (Fig. 5c, brown, compare top vs. bottom), with information decreasing as 1*/d*^2^ (cf. scaling of visual information as 1*/d*^4^, Eq. 1). Thus, even for target locations far from the starting wall, self motion cues are still sufficiently strong when the subject reaches them (compared to visual cues of the opposite wall). As a result, BION’s homing responses remain tethered to the starting wall (Fig. 5c, green and orange distributions; Fig. 5b, right), and we predict that this would remain the case even in more extremely expanded environments (Extended Data Fig. 10).

Just as human homing responses, rat grid fields were also shown to be tethered to environmental boundaries following unexpected environmental scaling^7^. In particular, the distance of a given grid field to the wall that the animal most recently approached (the equivalent of the ‘starting wall’ in human experiments) is preserved between familiar and compressed test environments (Fig. 5d, top left). In order for the neural posterior to follow the overall tethering of the ideal observer’s posterior to the last wall approached, which was apparent in the behavioral data (Fig. 5b, right), BION also predicted this tethering of neural tuning curves accurately (Fig. 5d, top right; Extended Data Fig. 11; r=0.98). Tethering along the changed axis was significantly larger in the compressed than in the familiar environment (both before and after testing in the compressed environment), and it was also larger than along the unchanged axis of the compressed environment (Fig. 5d, middle). Finally, tuning curves were significantly better explained by tethering than by scaling with the environment, both in experiments and in the model (Fig. 5d, bottom).

## Discussion

BION starts from the fundamental principle that representing spatial uncertainty is key for efficient and reliable navigation – and derives the pattern of homing responses and the deformation of grid fields from this principle. In contrast, most previous models did not consider spatial uncertainty and only explained a subset of the phenomena that we predict, and typically accounted for behavioral effects as mere epiphenomena of particular neural representations that were themselves often posited to be based on idiosyncratic features of the environment (such as ‘boundary vector cells’)^1,3,4,7,12,18,22,28–30,33–36^. For example, while successor representation-based theories explain some aspects of homing responses and grid field deformations^1,18,33^, their predictions about grid field diameters (or equivalently, radial frequencies, Fig. 1j in Ref. 1) deviate from experimental data^5^ and BION’s predictions (Fig. 2f), and they also do not explain experimental findings when the environment is changed (Figs. 3–5).

Models that did formalize navigation in a Bayesian inference framework have either only considered uncertainty about the identity of the environment, but not about location within the environment^37,38^, or they did not incorporate an image-computable ideal observer^16,21,37–44^. Thus, these models did not account for how environmental geometry affects spatial uncertainty and, in turn, homing behavior or grid cell responses.

Our work focuses on how a probabilistic estimate of location (and heading) is computed from noisy and ambiguous sensory inputs. A complete understanding of navigational behavior also requires the formalization of how these estimates, and the uncertainty about them, are used for route planning^45^ – something that we only modeled phenomenologically (Fig. 1c). Indeed, recent work using a more formal model of the planning process^16^ has been able to account for apparent violations of the Bayes-optimal combination of landmark and self-motion cues^46,47^ and reconcile them with earlier results suggesting optimal cue combination^14^. These results lend further support to our core suggestion: that Bayesian principles play a fundamental role in navigation.

## Acknowledgements

We thank T. Durrant for technical help with digitizing heatmaps. This work was supported by the National Research Foundation of Korea (NRF; RS-2025-16066185 and RS-2022-NR068758 to Y.K.) and KAIST (the International Joint Research Project to Y.K.) funded by the Ministry of Science and ICT of the Republic of Korea, the National Institutes of Health (R01NS117699 to D.M.W), the Air Force Office of Scientific Research (FA9550-22-1-0337 to D.M.W), the Wellcome Trust (Investigator Award in Science 212262/Z/18/Z to M.L.), and the Human Frontiers Science Programme (Research Grant RGP0044/2018 to M.L.). For the purpose of open access, the authors have applied a CC-BY public copyright license to any author accepted manuscript version arising from this submission.

## Methods

### Choice of experimental tasks to model

We chose experimental paradigms from the existing literature for which both human behavioral and rat grid cell data existed under analogous manipulations of environmental geometry (Table 1). Human behavioral data consisted of measures of endpoint distributions in homing tasks, while rat grid cell data consisted of geometric measures of grid fields (recorded while animals chased randomly dropped pellets).

**Table 1.** Experiments modeled in the paper.

| manipulation of environmental geometry | behavioral data (human) | neural data (rat) |
| --- | --- | --- |
| square vs. trapezoid | Bellmund et al. 2020 <sup>1</sup> | Krupic et al. 2015 <sup>5</sup> |
| expanded & compressed rectangles | Chen et al. 2015 <sup>3</sup><br>Hartley et al. 2004 <sup>2</sup><br>Keinath et al. 2021 <sup>4</sup> | Barry et al. 2007 <sup>6</sup><br>Keinath et al. 2018 <sup>7</sup> |

### BION model

We develop the ‘Bayesian image-computable observer for navigation’ (BION) model in which an agent infers its current allocentric pose (location and heading direction) based on a sequence of egocentric sensory observations. For this, we start from a generative model that formalizes the agent’s assumptions about the statistical relationship between its pose and sensory observations given its knowledge of the geometric layout of its environment. The agent then uses Bayesian inference to compute, and continually update, their posterior distribution over pose as new sensory observations arrive. The posterior expresses the probability with which the agent currently believes themselves to be in any given pose given past and current sensory information. In turn, these posterior distributions determine both behavior and grid cell responses.

#### Generative model

We consider an agent (a human or a rat) navigating in an environment, ℰ. In the experiments that we model, participants were extensively familiarized with the environment, or believed that they were in a familiar environment, so we make the simplifying assumption that the agent has complete (and perfect) knowledge of the configuration of the environment (also known as the ‘map’). Given a map, the agent uses a generative model to formalize its assumptions about the statistical relationships between its actions, its sensory inputs and its location, ℓ, and heading direction, *θ* (together, its pose: **z** = (ℓ, *θ*)). Importantly, as we model a wellcalibrated ideal observer, the same generative model also describes how the actual sensory inputs of the observer are generated (see also graphical model in Fig. 1a, and numerical parameter values in Table 2).

**Table 2.**
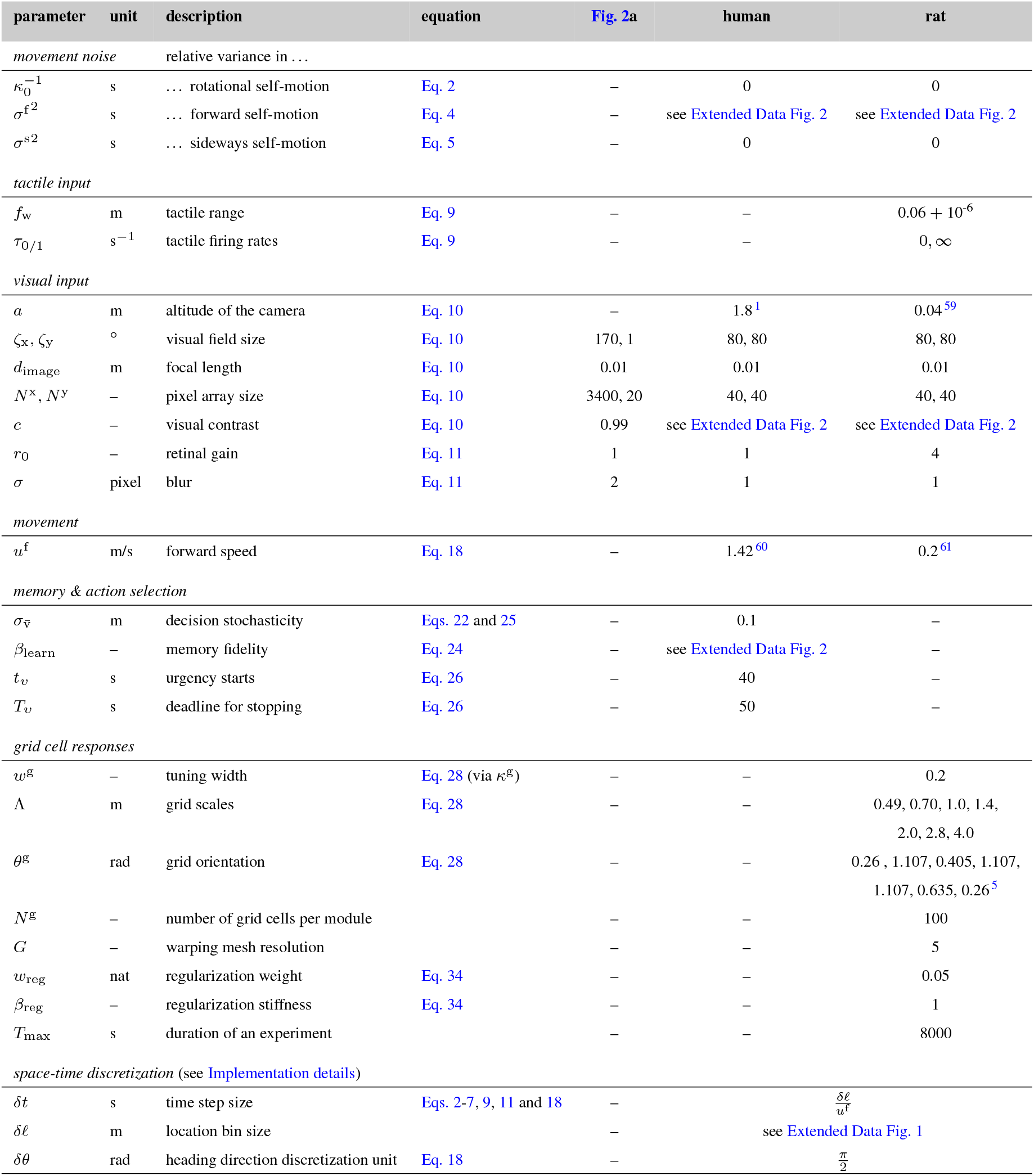
Model parameters. We used three distinct sets of parameters (last 3 columns). One set was used for generating Fig. 2a, which shows simulations of BION in a highly simplified setting (so that they can be compared with theoretical predictions, Eq. 1), only using visual inputs given a single, vertical edge extending across the height of the visual field (see also Methods, Comparison of analytical results with numerical simulation). Thus, in this case, parameters for other aspects of the model were irrelevant, as was the altitude of the camera. Human parameters were used for simulations shown in Fig. 1c, Fig. 2d, Fig. 3b-d, f, Fig. 5b-c, Extended Data Figs. 7, 8 and 10. Rat parameters were used for simulations shown in Fig. 1b, d-e (except that *r*_0_ = 2.7 was used here for illustrative purposes), Fig. 2b, e-f, Figs. 4 and 5d, Extended Data Figs. 3 and 4, Extended Data Fig. 5b, Extended Data Figs. 6, 9 and 11. Grid orientations were sampled with replacement from the set of experimentally observed values (Extended Data Table 1 of Ref. 5). Note that for each species, all but two (rat) or three (human) parameters were identical across all simulations. Values for experiment-specific parameters (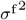, *c*, and *β*_learn_) are reported in Extended Data Fig. 2.

The agent takes actions and receives sensory inputs. We conceptually divide these inputs into self-motion inputs (primarily from the proprioceptive and vestibular systems), tactile inputs from whiskers (for rats) and visual inputs. Self-motion inputs inform the agent about the movements it makes in space, i.e. changes in its pose. We model these in a simplified way. Specifically, following Ref. 48, at each time step *t*, the agent’s actual change in heading direction (i.e. rotation), Δ*θ*_*t*_, is noisily related to its intended (and, for simplicity, perceived) rotational velocity,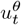:

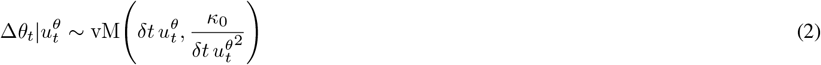

where δ*t* is the duration of a (discretized) time step, vM(·) is the von Mises (circular Gaussian) distribution (parameterized by its mean and concentration), describing the noisy relationship between perceived and actual rotation with signal-dependent motor noise^49^ (hence the inverse scaling of the concentration with 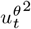), *κ*_0_ is its concentration for unit time and rotation, and 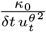 is sufficiently large (even for the maximal allowed rotational velocity 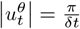) that the von Mises is well approximated by a normal distribution with a variance, 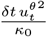, that scales linearly with δ*t* and 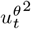, as appropriate. The agent’s updated heading direction is then simply:

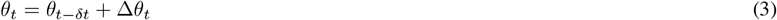

Similarly, also following Ref. 48, the agent’s actual translational movement (in egocentric coordinates), 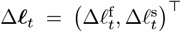, consisting of a forward, 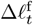, and a sideways component, 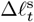, is also noisily related to its intended translational velocity, 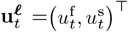. For simplicity, we only model intended translation in the forward direction (along the heading direction), so that 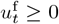 and 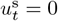. The actual forward movement, 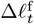 depends on the intended forward velocity, 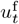, as:

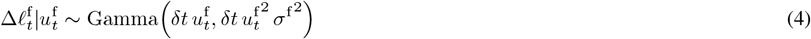

where Gamma(·) is the Gamma distribution (parameterized by its mean and variance) describing the noisy relationship between intended and actual forward movement, with signal-dependent motor noise^49^ (hence the scaling of the variance with 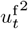) that also scales linearly with δ*t*, as appropriate. Although we only model intended translation in the forward direction (see above), we allow it to also lead to (unintended) sideways motion, with noise independent from that in forward movements, so that:

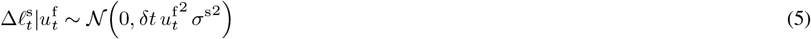

where 𝒩 (·) is the normal distribution describing this noisy relationship, again with signal-dependent noise (hence the scaling of the variance with 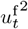) and scaling with δ*t*, as appropriate. The egocentric movement, Δℓ_t_, implies translation in allocentric coordinates that also depends on the heading direction of the animal (along which forward movements happen):

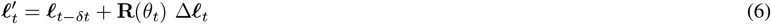

where 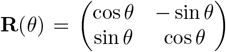 is the rotation matrix for angle *θ* and 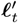 is the hypothetical next location in the absence of any boundaries. The actual translation of the agent is determined by one last factor: environmental boundaries, which cannot be crossed. We model this by limiting the agent’s actual translation to the farthest point along the line connecting its current, ℓ_*t*−1_, and the hypothetical next location, 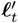 (Eq. 6), that it can still access without crossing an environmental boundary. Formally, the actual next location is given as

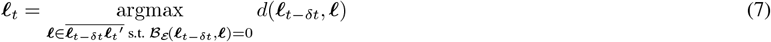

where 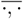 is the set of locations along the straight line connecting two locations, *d*(·, ·) is the (Euclidean) distance between two locations, and

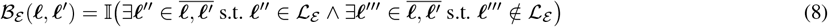

is an indicator variable for whether the line connecting two locations crosses an environmental boundary, with ℒ _ℰ_ denoting the set of locations that are inside the environment, ℰ.

Note that a recent study described a slowness prior that assigns higher probability to slower translation speeds and thus leads to undershoot in homing behavior, at least when self-motion inputs are restricted to optic flow^43^. We do not include such a slowness prior in the model because (1) in the experiments we modeled, vestibular inputs and efferent copies of motor signals were also available to participants (in addition to optic flow), allowing for more reliable perception of self-motion than in the experiments of Ref. 43; (2) our main results concern differences between conditions (e.g. differences in biases between expansion vs. compression of environments) rather than biases relative to true distances, such that any consistent, condition-independent bias would cancel out in the comparison.

Beside self-motion inputs, which are related to changes in its pose, the agent also receives sensory inputs that depend on its current pose. In particular, when simulating experiments in which rats navigated physical environments (but not for humans navigating virtual environments), we considered tactile inputs **w**_*t*_ = {*w*_*t,γ*_} from a set of sensors directed at different egocentric directions (relative to its heading direction), 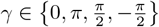 (i.e. to the front, back, left, and right side). Each of these tactile sensors (which could include whiskers as well as the tail, or more generally the surface of the body) is modeled simply as a Poisson ‘neuron’ whose rate depends on whether there is an environmental boundary (a physical wall in the rat experiments we modeled) within distance *f*_w_ in its corresponding direction:

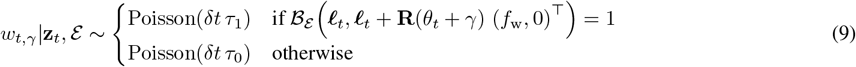

where *τ*_1*/*0_ is the firing rate of the neuron when the animal touches/does not touch a wall.

Finally, the observer also receives egocentric visual input from the environment. For this, the visual appearance of each environment, in particular with respect to its physical dimensions, was matched to that used in the corresponding experiment (Extended Data Fig. 1). In the original experiments, distal landmarks beyond the experimental arena provided cues that could be used to disambiguate the otherwise identical-looking walls of the arena. In lieu of such distal landmarks, we rendered each wall in a unique color to make walls identifiable.

Visual input consists of the activation of a flat array of retinotopic neurons for each color channel, *c*_p_ ∈ {R, G, B}, determined by an idealized pinhole projection of the 3D scene:

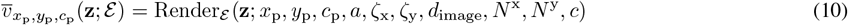

where 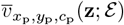 is the value of the pixel at position (*x*_p_, *y*_p_) in the imaging plane, when color channel *c*_p_ of environment ℰ is rendered through a pinhole camera (with an infinitesimal aperture) from a vantage point defined by pose **z**_*t*_ = (ℓ_*t*_, *θ*_*t*_) (i.e. the camera is placed at 2D location ℓ_*t*_ with its optical axis directed at direction *θ*_*t*_ in the horizontal plane). The camera is at (constant) altitude *a* and further characterized by its horizontal and vertical visual field size, ζ_x_ and ζ_y_, respectively, and the distance of the imaging plane from the aperture, *d*_image_. The imaging plane is a flat rectangle, with a pair of horizontal and a pair of vertical sides, such that it is orthogonal to and centered on the optical axis, with pixel positions defined on an *N* ^x^ × *N* ^y^ regular grid, such that *x*_p_ and *y*_p_ take regularly spaced values on the ±*d*_image_ tan 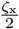 and ±*d*_image_ tan 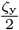 intervals along the horizontal and vertical sides, respectively. Visual input was rendered at contrast *c*. (As *c* scales the rendered pixel values linearly, and neural responses are in turn linearly related to pixels, Eq. 11, changing *c* effectively implies a scaling of neural tuning curves.) Render _ℰ_ (·) was computed by using the Mayavi package^50^ (version 4.7.4).

Visual neuron activations are given by **v**_*t*_, such that neuron *i* integrates pixel values around position (*x*_p_(*i*), *y*_p_(*i*)) in the grid for color channel *c*_p_(*i*), using a Gaussian receptive field, with its response being corrupted by Poisson noise:

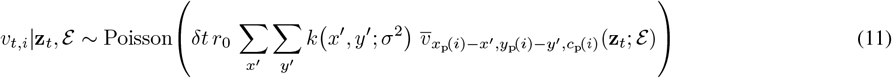

where, for these purposes, 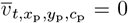 for pixel locations (*x*_p_, *y*_p_) beyond the grid, *k*(*x*^*′*^, *y*^*′*^; *σ*) is a (zero-mean) Gaussian with (spherical) variance *σ*^2^, truncated at ±3 *σ* along each dimension, and renormalized so that ∑_*x*_*′* ∑_*y*_*′ k* (*x*^*′*^, *y*^*′*^; *σ*^2^)= 1, and *r*_0_ is a scaling factor expressing the change in visual neuron firing rate for a unit change in pixel values.

Although this model is more realistic than previous models that assumed direct bearing and distance information about predefined landmarks^16^, it is still obviously a highly simplified model of visual inputs. In particular, it only considers a single, ‘cyclopean’, frontally looking eye with infinite depth of field, and a flat (rather than curved) retina with spatially homogeneous resolution in the plane (with simple Gaussian blur) that does not depend on eccentricity. Moreover, for simplicity, we set the parameters (apart from *a*) of this part of the model in a species-independent manner and with values that very approximately (if at all) matched the actual parameters of rat and human eyes. While these simplifications may change the quantitative details of our model predictions, we do not expect them to affect our results qualitatively. For example, with binocular vision, visual disparity provides additional information about depth. However, a mathematical analysis suggests that beyond an overall scaling effect (which can be subsumed into our contrast parameter, *c*), binocular vision only leads to additional *<* 1.5% (for rats) or *<* 0.2% (for humans) pose-dependent modulation of visual information in the experiments we considered (Supp. Material SM1).

#### The ideal observer

The ideal observer (Fig. 1b) is given by the Bayesian inversion of the generative model. Specifically, in each time step *t*, it computes the posterior over the current pose, **z**_*t*_ = (ℓ_*t*_, *θ*_*t*_) (the combination of location ℓ_*t*_ and heading direction *θ*_*t*_), given the history of sensory inputs up to (and including) the current time step, **s**_1:*t*_ (where 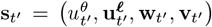). This posterior, 𝒫^ideal^(**z**_*t*_|**s**_1:*t*_), expresses the probability with which the ideal observer believes its current pose to be **z**_*t*_. Importantly, it can be computed by recursive Bayesian filtering, i.e. by updating the posterior computed in the previous time step with only the sensory input received in the current time step (so that sensory inputs in previous time steps do not need to be memorized):

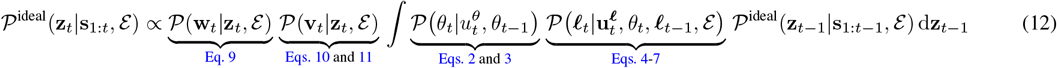

where the four terms needed for updating the posterior are given by the equations defining the generative model as indicated, with 𝒫^ideal^(**z**_*t*−1_ | **s**_1:*t*−1_, ℰ) replaced by the prior 𝒫^ideal^(**z**_0_ |ℰ) for *t* = 1 (see also below). In particular, as Eqs. 10 and 11 define 𝒫 (**v**_*t*_ |**z**_*t*_) by a generative model for a fully rendered visual image, this ideal observer is *image-computable*, i.e. it can use any visual image as input for inferring pose, without having to provide explicit information about its current pose with respect to environmental landmarks or boundaries. Note that we made the dependence of the individual terms in Eq. 12 on the environment, ℰ, explicit. (The term describing self-motion inputs about rotations is an exception because these do not normally depend on the current environment.) While, by construction, sensory inputs always depend on the actual environment in which the agent is actually navigating (ℰ ^∗^), the agent conditions its inferences on the environment it believes it is navigating (ℰ). Critically, as we explain below, in some experiments, there may be a model mismatch, i.e. the believed environment may not match the actual environment (ℰ≠ ℰ ^∗^). The posterior inferred by the ideal observer, and in particular, the marginal posterior over locations

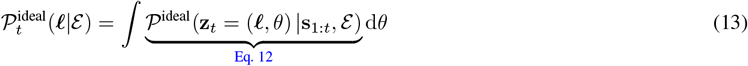

then forms the basis of the model’s predictions for behavioral (homing) and neural (grid cell) responses.

To initialize the Bayesian filtering algorithm (Eq. 12), we use a prior to describe the observer’s belief about its pose in the ‘zeroth’ time step, before it has received any sensory input, 𝒫^ideal^(**z**_0_|ℰ). See Priors for experiment-specific choices of priors.

#### Behavioral responses: cue-driven journeys

In order to model behavior, i.e. to determine the sequence of control signals *u*^*θ*^ and **u**^ℓ^ (see Generative model above), we distinguish between cases when participants needed to follow explicit sensory cues as to where they should go in an environment (cue-driven journeys), and when they needed to rely on memory to decide where they wanted to go (memory-based journeys). All experiments with rats that we model used a random pellet-chasing paradigm and so we considered rats’ journeys to be purely cue-driven. For human experiments, the learning phase consisted of cue-driven journeys towards visible targets (objects), while the test phase consisted of memory-driven journeys where participants needed to navigate towards previously seen target locations based on their memory of those locations (in some cases, these memory-driven journeys were preceded by a cue-driven journey for ‘initialization’, see below). For some experiments, the learning and test phases were conducted in separate trials^1,2,7^, while for Ref. 3 each trial started with a learning phase and ended with a test phase.

After initialization (see Experiment-specific simulation details), we model behavior as greedy, but noisy, minimization of the distance between the observer’s current (inferred) location and a target location (determined by either a sensory cue controlled by the experimenter, or by a memory of a location stored by the observer). Specifically, in each time step *t*, the model considers a set of *C* + 1 distinct ‘control signals’ (i.e. ways of changing its pose):

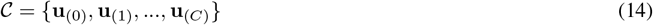

where each control signal consists of a combination of intended rotation and translation:

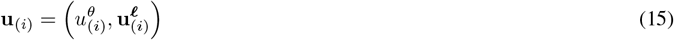

with translation itself being a combination of forward and sideways component, the latter being fixed at 0 (see also Generative model)

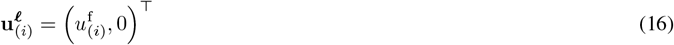

Specifically,

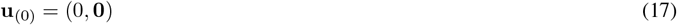

is the ‘stop moving’ action, and to simplify route planning, for *i >* 0, we use

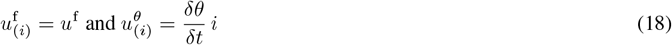

where *u*^f^ is a constant forward motion speed, δ*θ* is the unit of discretization used for heading direction, δ*t* is the temporal bin size used in a given simulation (see Implementation details and Table 2), and 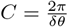 (integer with our choice of δ*θ*; Table 2).

For route planning, the model simulates the counterfactual consequences of executing each control signal in terms of how it would change (the posterior over) its location in the next time step (cf. Eq. 12):

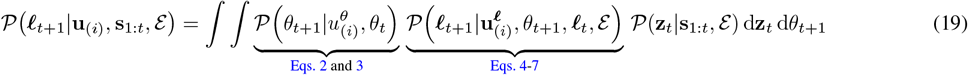

where 𝒫 (**z**_*t*_|**s**_1:*t*_, ℰ) is the observer’s posterior over its current pose, which for cue-driven journeys (during which experimental participants tend to move systematically towards the cue) was simply chosen to be concentrated on its veridical pose: 𝒫 (**z**_*t*_|**s**_1:*t*_, ℰ) = δ(**z**_*t*_ − **z**^∗^_*t*_). These counterfactual posteriors are then used to compute the expected value of each control signal:

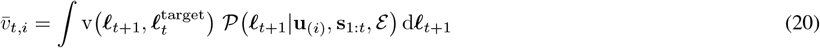

where 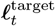 is the current target location, and the value of being at location ℓ when the target location is ℓ^target^ is defined to be the negative squared Euclidean distance between the two locations:

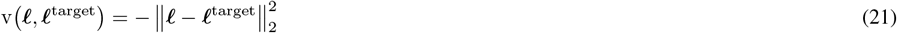

and the next control signal is chosen using a softmax decision rule based on these expected values:

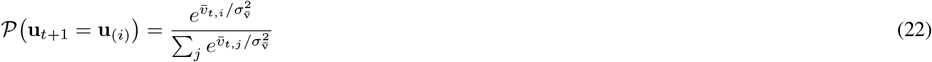

with decision stochasticity 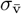 (effectively scaling the distance in Eq. 21, such that 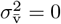 corresponds to deterministically choosing the control signal with the highest value, and 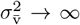corresponds to choosing control signals with uniform probabilities). This results in a myopic policy locally (and stochastically) minimizing Euclidean distance to the target location.

For rat experiments, target locations (corresponding to randomly dropped pellets) are chosen sequentially to maximize coverage. Specifically, at the beginning of a trial, or once the agent arrives at the currently cued location (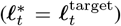), a target location is chosen randomly among those locations that would maximize the minimum number of visits to a spatial bin, assuming the agent would follow a straight path between its current location and each candidate new target location. This target location persists until the agent arrives at it. See below for human experiments (distance estimation task).

#### Predictions for memory-based journeys: homing task

Homing experiments consist of a learning and a test phase. In the learning phase, participants memorize the location of an object (or the locations of several visually distinct objects) to which they are made to navigate along cue-driven journeys (see above). We model the memory of such a ‘target location’ as the mean location under the ideal observer’s posterior (Eq. 13) at the time, *T*_learn_, it reaches the corresponding object (conditioned on the stimuli it received on its journey up to that point) after memory decay:

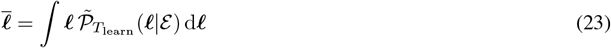

where

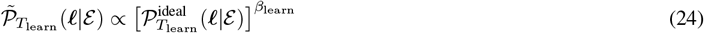

and 0 *< β*_learn_ ≤ 1 is the inverse temperature parameter controlling the level of memory decay (e.g. due to forgetting; the lower, the more decay).

In the test phase, we model the participant’s behavior as we do for cue-driven journeys, except that counterfactual next-step posteriors are computed using the ideal observers posterior over its current location (𝒫 (**z**_*t*_|**s**_1:*t*_, ℰ) = 𝒫^ideal^(**z**_*t*_|**s**_1:*t*_, ℰ) in Eq. 19), and the target location throughout a given test trial is the memorized location of the object cued in that trial (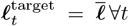 in Eq. 20, see Experiment-specific simulation details for modeling the experiments in Ref. 4 that involved multiple learning and testing phases for the target location.) In addition, action selection also includes an urgency signal prioritizing stopping as time passes, such that:

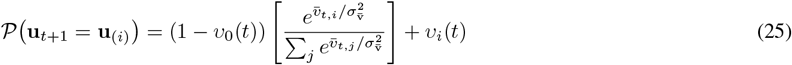

(cf. Eq. 22), where the urgency signal encourages the choice of stopping after *t*_υ_, with linearly increasing propensity until reaching *T*_υ_, when the choice of stopping is deterministically forced^51–53^:

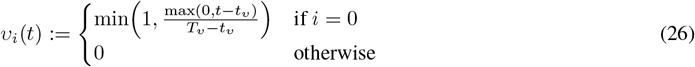

(The urgency signal made no discernible contributions to the predicted routes in most cases, except when the true environment was compressed and the agent moved along the compressed axis. In such cases, the urgency signal prevented an infinite loop whereby the agent would start believing that the goal is behind it as soon as it approaches either wall at the end of the compressed axis, which necessarily looks closer than expected.)

The route is terminated when the agent chooses the special ‘stop moving’ control signal **u**_*t*+1_ = **u**_(0)_ (Eq. 17), and its true location at that time, ℓ_*t*_, is recorded as the responded location.

For every experiment considered, the number of simulations is (approximately) matched to the corresponding original data, in terms of the number of subjects and the number of trials per conditions and subjects (Extended Data Fig. 1). We analyze these homing responses following the same procedures used for the original analyses in the corresponding experiments (see also Experiment-specific simulation details).

#### Predictions for grid cell responses

Here we describe a decoding-based approach for relating grid cell responses to the ideal observer’s posterior, which can be most directly related to standard approaches to decoding the animal’s location from grid cell responses^54^. In the Supplementary Material (Supp. Material SM2) we describe an alternative, encoding-based approach, in which grid cell responses are defined directly as functionals of the ideal observer posterior, and show that it gives near-identical results without assuming that cells have access to the true location of the animals, the last wall it approached, or the identity of its actual (rather than believed) environment.

We model a population of grid cells, grouped into discrete modules (see below), such that the firing rate, *r*_*t,i*_, of cell *i* at time *t* is defined by its tuning curve and the current ‘state’ of the animal:

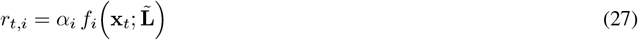

where *α*_*i*_ is a gain parameter scaling the tuning curve of the neuron (shared by cells in a module), **x**_*t*_ = (ℓ_*t*_, *ω*_*t*_) is the state of the animal relevant for characterizing tuning curves, combining (as described experimentally^7,55^) its 2-dimensional location in the environment, ℓ_*t*_, and a categorical variable indicating the last wall it approached (defined as coming within 12 cm of that wall^7^), *ω*_*t*_ ∈ {west, east, north, south}, and 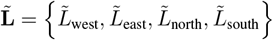 is a set of ‘warping’ parameters (shared by all cells) controlling the shapes of the grid cell tuning curves in an *ω*-dependent manner (see below).

Each grid cell tuning curve has multiple location-tuned grid fields arranged on a hexagonal lattice, obtained (following earlier approaches^5^) as a product of three planar waves at regular 120^°^ angles, but such that the lattice is warped based on 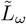:

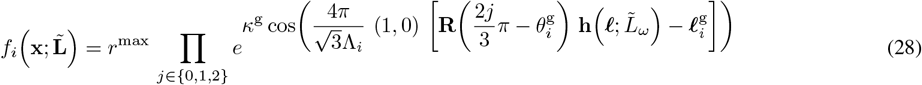

where *r*^max^ and 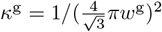 (where *w*^g^ is approximately proportional to the width of the grid field relative to the grid scale) are the maximal firing rate and the tuning precision of the neuron (shared across all grid cells), and Λ_*i*_, 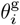, and 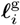 are the grid scale (the distance between neighboring peaks), orientation, and preferred phase of the neuron (different across individual grid cells or grid cell modules, see below). Critically, **h**(·) denotes (inverse) warping, such that grid fields would be regular in the unwarped space. The warping is defined as a bilinear interpolation (implemented by PyTorch’s grid_sample function^56^) between the *G× G* control points of a (*ω*-dependent) mesh, 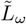.

The control points parameterizing the warping, 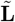, are shared across all neurons. (Although module-specific distortions of grid fields have been described^57^, these have been explained away by tethering effects^7^ that our model also captures, Fig. 5, and so we do not consider making 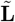 different across modules.) Warping parameters are optimized separately for each experimental paradigm as appropriate (see below). All other (unwarped) grid cell parameters are fixed across experiments and chosen following experimental work on grid field geometries in undeformed environments (see also Table 2). Specifically, neurons are grouped into seven discrete modules^57^, with the *N* ^g^ cells of a given module sharing the same grid scale, Λ_*i*_, and orientation, 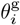, but differing in their preferred phases, 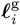. Module-specific grid scales are chosen to be a geometric series with a 1.42 quotient^57^, while grid orientations (relative to a horizontal wall) for each module are sampled uniformly with replacement from a set of experimentally described values^5^. Cell-specific preferred phases, 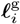, are uniformly sampled from an equilateral triangle with sides Λ_*i*_ to ensure a uniform coverage of space (on average) by the grid fields of cells corresponding to each module^55^. Finally, maximal firing rates, *r*^max^, and tuning precision, *κ*^g^, are shared across all neurons and set to values that provide realistic tuning curves.

Grid field warping is optimized such that a probabilistic decoder of grid cell responses produces a posterior, the ‘neural posterior’, that maximally matches the posterior of the ideal observer (Eq. 13) at any time. Assuming a large population of grid cells with independent Poisson noise in their firing, the neural posterior is decoded from grid cell responses using a product-of-experts decoder (such as that employed by probabilistic population codes^24^):

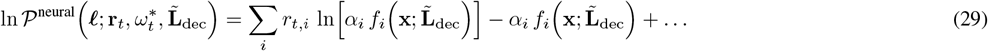

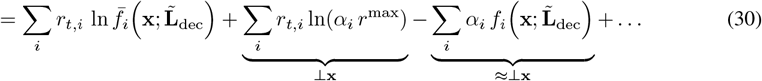

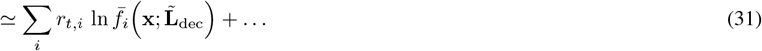

where ℓ is (a candidate) decoded position, **r**_*t*_ is the firing rates of the grid cell population at time *t*, 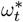 is the identity of the last wall approached (which is thus assumed to be known to the decoder), 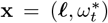 is the combination of the decoded position and the identity of the last wall approached, 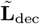 is the warping parameters of the ‘decoding’ tuning curves (see Eqs. 27 and 28),0020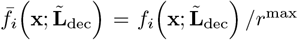 are the normalized decoding tuning curves, ⊥ **x** and ≈⊥ **x** respectively denote terms that are exactly or approximately independent of ℓ, and as such are ignored in the following, and … denotes such terms once they have been ignored. (Note that we can treat the last term in Eq. 30 as approximately constant with respect to **x** following standard approaches^58^, as tuning curves are sufficiently evenly and densely packed before warping, and the unevenness introduced by warping does not affect this constancy. As a consequence, the neural posterior does not depend on the gains of the decoding tuning curves – only on the gains of the encoding tuning curves through **r**_*t*_.)

In order to understand how optimization proceeds, it is useful to rewrite Eq. 31 such that its dependence on the actual state of the animal, 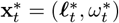, determining the firing rates of grid cells, 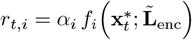, based on the gains and warping parameters of the ‘encoding’ tuning curves, *α*_*i*_ and 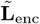 (Eqs. 27 and 28) becomes explicit:

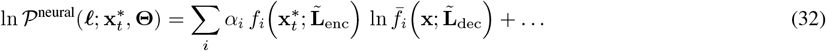

where, in order to simplify notation, we introduced 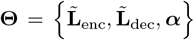 to denote the collection of all optimizable neural parameters. Critically, the encoding and decoding tuning curves may not be the same^3^: the encoding tuning curves are defined in the actual environment the animal is navigating, ℰ^∗^, while decoding tuning curves are defined in the environment in which it believes to be located, ℰ. Just as for the ideal observer above, we distinguish between ℰ^∗^ and ℰ because in some experiments the two were not the same: the animal had been familiarized in one environment, and then unbeknownst to them was tested in a different (scaled) environment (Extended Data Fig. 1 and figs. 3 and 5). In such cases, we take the actual environment to be the test environment, while the believed environment to be the familiar environment.

We quantify the (mis)match between the ideal observer’s and the neural posterior (Eqs. 13 and 32) via the average moment-by-moment Kullback-Leibler divergence (or, in this case, equivalently, the cross-entropy) between them (note that the two posteriors are computed over the same ‘believed’ environment, ℰ, as appropriate):

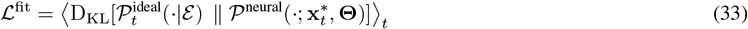

We optimize the warping to minimize the measure of mismatch in Eq. 33 together with a regularizer that penalizes deviation of the angle between neighboring nodes of the mesh from 90^°^:

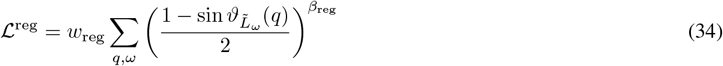

where *q* runs through all triads of control points that define a corner of a cell in the mesh, 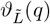 is the angle of such a corner in a warped mesh 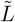, and *w*_reg_ and *β*_reg_ are hyperparameters that control the weight and stiffness of this regularization, respectively. Therefore, the total loss we optimize is

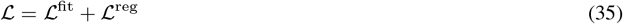

with the constraint that 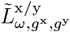 is monotonic in *g*^x*/*y^ ∀ω, g^y*/*x^ to further reduce entanglement of the warped mesh.

### Parameter choices

For most parameters, we used a single set of values for modeling all human experiments, and another set for modeling all rat experiments (Table 2). Also for most parameters, we chose values to have a reasonable match with experimental data (e.g. forward speed, *u*^f^). Some parameters were set to extreme values for simplicity: rotational inverse variance *κ*_0_ was set to be large enough to make rotations deterministic (as the experiments we simulated had orientation cues), and similarly, sideways motion noise variance *σ*^s^ was set to zero, and tactile firing rates to *τ*_0_ = 0 and *τ*_1_→∞ (which make tactile inputs noiseless, while still retaining ambiguity, as different poses can result in the same tactile input). We minimized the discrepancy between model predictions and experimental results by very coarsely optimizing two or three model parameters (for rat and human experiments, respectively). For this, we performed a grid search with only a few possible values for each parameter (Extended Data Fig. 2): contrast (*c*, 3 values), and relative variance in forward self-motion (*σ*^f^, 3 values) for both human and rat experiments, and memory fidelity (*β*_learn_, 2 values) for human experiments. In addition, for rat experiments, we similarly optimized the scale of analyzed grid cells (5 values, see Comparing modeled and experimental grid cell responses below). Only such a coarse optimization was possible due to the excessive computational cost of simulating the model for any combination of parameters. The objective of optimization was to minimize the sum of discrepancies across all comparisons we sought to reproduce in the given experiment (Extended Data Fig. 2). To measure discrepancy between model and experiments for a given comparison, we performed the same statistical comparison on simulated data as in the original experiments (see Analyses of simulation results), and used a metric that depended on whether simulations matched the sign and significance of the experimental comparison (but not on matching its magnitude; Extended Data Fig. 2a).

### Experiment-specific simulation details

While we used the same model to simulate all experiments, we allowed a minimal number of parameters to be different across human and rat experiments to capture some critical differences between the two species (see above, Parameter choices), and also took into account differences in the details of the specific experimental paradigms that we simulated (see also Extended Data Fig. 1 for the environmental layouts used in different experiments).

#### Priors

For modeling rat experiments (Fig. 2e & f, Fig. 3f & g, Fig. 5d), we use a uniform prior over all poses, i.e. uniform distribution over heading directions, and a uniform distribution over locations within the boundaries of the believed environment, ℰ. The precise choice of the prior in this case has little practical relevance, because we run long trials, with a long sequence of observations (26766 time steps), and then discard the initial time steps (100), so that the analyzed portion of our simulations has negligible dependence on the prior). For modeling human experiments in which participants were consistently started from a small subset of initial poses across trials (Fig. 3a-e, Fig. 5a & b), we match the prior to the true generative process, i.e. to the distribution that was actually used to initialize participants’ poses in the corresponding experiments (see also Behavioral responses: cue-driven journeys). For modeling the experiment in Ref. 1 (Fig. 2c & d), which used a large set of potential initial poses generated by a complicated set of rules that was unlikely to have been incorporated into participants’ priors, we use a uniform distribution over all possible poses as the prior.

#### Initial poses

For modeling rat experiments (Fig. 2e & f, Fig. 3f & g, Fig. 5d), we sample the initial pose of the agent from a uniform distribution, to be consistent with our prior (although, the choice of this distribution had minimal relevance, see also above The ideal observer). For modeling human experiments, we match the distribution of starting poses to those used in the corresponding experiments (chosen using the same procedure across all trials of an experiment, even if they corresponded to separate learning and test phases, as in Refs. 1,2,7, see above). Specifically, for modeling Ref. 1 (Fig. 2c & d), we sampled the starting location of the agent such that (1) it was at least 12.5 m away from the true target location, (2) in half of the trials, it was in the same half of the environment as the target location and in the other half of the trials it was in the opposite half, (3) its distance to the target position was matched between conditions (halves of the two environments; personal communication by Jacob Bellmund). For modeling Ref. 2 (Fig. 3a & b), the agent was randomly placed at one of the four corners of the environment and faced one of the two outward facing directions (p.48 of Ref. 2). For modeling Ref. 3 (Fig. 3c-e), the agent was always at the center of the environment and faced a random direction. For modeling Ref. 4 (Fig. 5a & b), the agent was placed at the center of one of the four walls, facing outward.

#### Multiple learning and test phases for the same target location

In the experiments reported in Ref. 4 each target location was learned in multiple learning phases (in their terms, ‘collect’ trials). Specifically, each target location was first learned in two learning trials, and then it was learned again in four pairs of test and learning phases (in that order). Subjects started each of the six learning phases from a random location, although they started each test phase facing the center of one of four walls in a random order. We modeled this process by computing the average of the memorized locations from each learning phase preceding the given test phase (computed as in Eq. 23). This is equivalent to maximizing the value v(ℓ, ℓ^target^) (Eq. 21) when ℓ^target^ can be one of previously memorized locations with equal probabilities.

#### Distance estimation task

In addition to the homing task, participants in Ref. 1 also performed a distance estimation task. In this task, during the test phase, participants were asked to estimate the distance between two target objects they had seen during the learning phase. Participants indicated their estimate by walking the corresponding distance in a virtual circular enclosure (to which they were previously familiarized). Specifically, they were asked to walk the same distance as the distance between a pair of objects, starting from the root of a static visual arrow placed randomly in the familiar circular environment, following the direction of the arrow, which always pointed towards the center of the enclosure.

In our simulation, the observer moves along a cue-driven journey from a random initial location to the starting point of its memory-based journey, corresponding to the root of the arrow in the original experiment (which we did not explicitly render in our simulations), and then rotates until it faces the center of the environment (i.e. at this point only, the set of control signals 𝒞, Eq. 14, also included pure rotations). Thus, the observer starts the memory-driven journey from this root position, with a distribution of believed locations 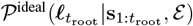. During the memory-based journey, behavior is modeled similar to the homing task with two key differences. First, only those control signals are considered that do not have a rotational component (see above) so that the observer keeps moving towards the center of the environment as in the original experiment. Second, the expected value of a control signal depends on how well the distance between the resulting next location and the root location matches the distance between the remembered locations of the two targets, 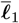 and 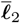, and so uncertainty about both the root and the (counterfactual) next location needs to be integrated out (cf. Eqs. 20 and 21):

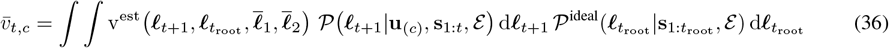

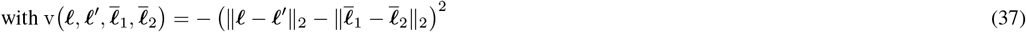

where 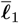 and 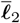 are computed as for the homing experiment (Eqs. 23 and 24). The observer’s true distance from the root location at the time when it chooses to stop is recorded as the responded distance.

#### Optimizing warping parameters for encoding vs. decoding grid fields

For modeling experiments using static environments^5^, in which neural responses are analyzed in the same environment with which the animal has been familiarized (Fig. 2e-f), 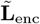 and 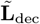 are tied (i.e. kept identical throughout optimization), and optimized jointly with ***α*** to minimize ℒ (Eq. 35). For modeling ‘scaling’ experiments^6,7^, in which neural responses are analyzed after an unexpected scaling of the familiar environment (Fig. 3f-g, and Fig. 5d), parameters are first optimized for encoding posteriors while the animal is moving in the familiar environment, as described above, following which 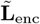 and 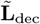 are untied, such that 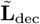 remains fixed (so that decoded positions are still expressed in the familiar environment, as appropriate given that animals are assumed not to be aware of the change in the environment), but 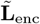 is re-optimized (jointly with ***α***) to minimize ℒ in the test environment (so that firing rates are defined by the animal’s location in the current test environment, to make it comparable with experimental data in which grid fields are mapped in the test environment by definition).

### Implementation details

All our simulations were discretized in time and space. The movements of the observer were simulated at a temporal resolution of δ*t*, and its pose was represented at a spatial resolution of δℓ for locations on a square grid and δ*θ* for heading directions (Table 2 and Extended Data Fig. 1). To simplify route planning, we chose δ*t* such that *u*^f^ δ*t* corresponded to one spatial bin in each environment. Location bin sizes δℓ were set to be small enough to reliably capture the effects of manipulations in the experiments simulated, and large enough to keep the demand on compute and memory to a reasonable level.

Distributions describing noise in actual movements relative to intended movements (as determined by the control signals) were discretized by evaluating the corresponding probability density functions (Eqs. 2, 4 and 5) on a grid with the same spatial resolution and then normalizing these values to obtain a well-defined discrete distribution. (Note that because our discretization only allowed cardinal heading directions, and possible boundary crossings were only evaluated at the same spatial resolution, translations always led to movements aligned with the simple square grid of spatial bins used for discretizing locations; Eqs. 6 and 7.)

To ensure that the ideal observer remained well calibrated, its inferences were computed on the same spatially discretized grid and at the same temporal resolution as its real movements, using the same discretized distributions (with integrals Eqs. 12, 13, 19, 20, 23 and 36 thus approximated by sums as appropriate). In experiments involving environmental scaling (Figs. 3 and 5), movements were simulated on the grid of the actual environments, whereas inferences were computed on the grid of the familiar environments.

Neural posteriors were also, by necessity, computed at the same spatial resolution as the ideal observer’s posterior, using the same discretization approach based on evaluating the continuous densities at the spatial resolution of our simulations and then normalizing as appropriate (Eq. 32). As a consequence, divergences between ideal observer and neural posteriors were computed as sums over the spatial grid of discretized locations (Eqs. 33, 38 and 42-44). In particular, to accelerate the computation of the average divergence in Eq. 33, which needed to be performed at every iteration of the procedure used for optimizing grid field warpings, we rewrote it as

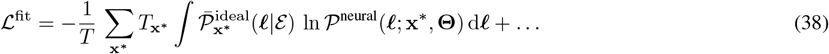

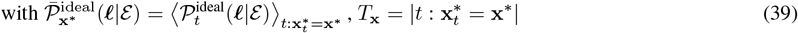

where **x**^∗^ indexes the discretized poses of the animal, … denotes additive terms that do not depend on **Θ**, and we note that 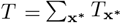. The quantities defined in Eq. 39 also do not depend on **Θ** and so can be pre-computed, which greatly accelerates the optimization of **Θ** when |{**x}**|**≪** T, as was the case for our simulations (and the integral in Eq. 38 was also approximated by a finite sum, as in the other cases described above). Optimization of the grid field warping was performed with gradient descent, by feeding the gradient computed with PyTorch 2.1.2 to the BFGS algorithm provided with the optimize.minimize function of scipy 1.13.1 with default settings.

For simulating each rat experiment, we simulated one session (trial) while simulating the activity of a large number of cells. As the original experiments were limited in the number of cells that could be recorded simultaneously, they used multiple sessions, and pooled recorded cells across sessions. Thus, rather than exactly matching the duration of individual sessions of the experiments (40^5^, 20^6^, and 15-30^7,57^ minutes), we simulated a single session with a 133.3 minutes (8000 seconds) duration to ensure a good coverage of poses while keeping the duration in the same order of magnitude as multiple sessions of an experiment.

### Analyses of simulation results

In general, to compare model simulations to experimental results, we used the same statistical tests on the same (or virtually the same) number of samples as the original experiments (see details for specific experiments below).

#### Goodness of fit

We quantified the goodness of fit of BION to experimental data for each result showing a direct model-experiment comparison of the variations of some summary statistics (bias, scaling, gridness, etc.) across conditions. (These results are presented as bar plots: Fig. 2d and f, Fig. 3d, Fig. 3f and Extended Data Fig. 8, Fig. 4b-c, Fig. 5d.) To measure how well BION reproduced the overall pattern of variations seen in the data for a given experiment, we computed *r* as the correlation between experimental results and corresponding model predictions (across all summary statistics and conditions reported for that experiment). Correlations were computed under an ordinary linear regression model with a separate pair of coefficients (offset and scale) for each summary statistic (excluding summary statistics for which only two conditions were analyzed; Fig. 5b, Extended Data Fig. 5b), to accommodate the potentially different magnitudes and units associated with different summary statistics. Note that the parameters of BION were not optimized for *r*, only for a more qualitative metric of model (mis)match (Extended Data Fig. 2), and even for that, only 2–3 parameters were optimized very coarsely (see Parameter choices above).

#### Comparison of analytical results with numerical simulation

For Fig. 2a, we compared the visual Fisher information given by the analytic solution Eq. 1 with numerical simulation. For the simulation, we used a single straight upright edge placed at various distance and eccentricity, formed with two walls stretching vertically to fill the visual field. The Fisher information was computed with the partial derivative of the tuning curve computed with the finite difference method.

#### Variation of visual information within an environment

For Fig. 2b, we quantified the agent’s visual uncertainty *σ*^x*/*y^ as the square root of the variance of the visual posterior along each axis x*/*y at each spatial bin ℓ^*′*^, averaged across the times when the agent was at that spatial bin:

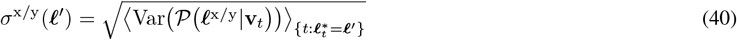

Similarly, we computed the uncertainty combined across axes as follows:

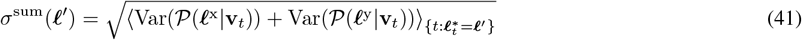

We also show a scale for the spatial information (on the same color bar as the uncertainty) as the inverse mean variance 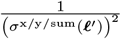 as the spatial information.

#### Analyses of homing responses and distance estimates

In general, for a direct comparison, we analyzed simulation results using the same methods that were used to analyze data in the corresponding experiments. Specifically, for Fig. 2d, top, we analyzed homing response locations following methods for Fig. 2b & e in Ref. 1. For Fig. 3b, we followed methods for Fig. 2 of Ref. 2. For Fig. 3d, we followed methods for Fig. 4A and Fig. S4 of Ref. 3, except that we subtracted the mean response of the baseline condition (‘matched’ return journey in an unchanged environment) within each experiment to reveal changes in homing locations due to the scaling of the environment and to reduce clutter in the figures. For a fair comparison, we did this both for plotting simulated data and for re-plotting experimental data from the original experiments for means. In the plot, error bars for the model show s.e.m. of relative biases across simulated participants. As experimental data were not available for individual participants, error bars for experiments show s.e.m. of absolute biases in the test environment (ignoring s.e.m. of data in the familiar environment), but statistical significances are shown for relative biases as in the original publication (see also Extended Data Fig. 8)^3^. For Fig. 5b, we followed methods for Fig. 6d of Ref. 4. For Fig. 2d, bottom, we analyzed homing response locations following methods for Fig. 3c & e in Ref. 1, except that we subtracted the mean estimated distance in square environments to reveal the differences between estimates. (Again, for a fair comparison, we did this both for plotting simulated data and for re-plotting experimental data from the original experiments.)

#### Characterizing the quality of optimized grid field warpings

In Fig. 1e, to assess the quality of the match achieved by the optimized neural parameters, **Θ**^∗^ (minimizing ℒ, Eq. 35, for the trapezoid environment as in Fig. 2e-f), we show the original (average) D_KL_ used for computing ℒ^fit^ (Eq. 33) (‘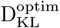 ‘, purple) along with three controls, which are defined as follows:

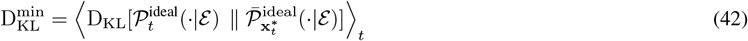

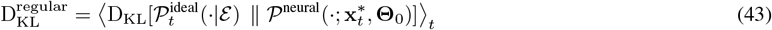

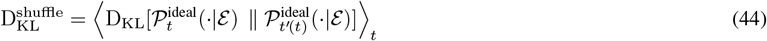

where **Θ**_0_ is the warping parameters that give regular grid fields (for both 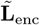 and 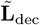), and *t*^*′*^(*t*) is the *t*-th time step in a permutation of time steps. Eq. 42 represents a lower bound on the divergence attainable between the ideal observer and neural posteriors taking into account the fact that the neural posterior was constrained to depend only the current location of the animal and the last wall it approached (Eq. 32), while the ideal observer posterior in general depends on the full history of observations (Eq. 12, see also Eq. 39). Eq. 43 provides calibration for the importance of warping by expressing the divergence attained without any warping. Finally, Eq. 44 represents an effective upper bound by expressing the divergence of the ideal observer posterior between two randomly selected time points.

#### Comparing modeled and experimental grid cell responses

For a comparison with experimental data, optimized grid cell tuning curves were computed (using Eq. 28) at a 2.5 cm resolution (Figs. 2, 4 and 5), matching the spatial bin sizes used for analyzing experimentally recorded grid cell tuning curves in the corresponding experiments^5–7^. Our analyses of grid cell tuning curves were based on grid cells whose scales after deformation was 24–55 cm for Ref. 5 and 30–130 cm for Refs. 6 and 7, matching the range of scales of experimentally recorded grid cells that underlay the analyses of grid field geometries. We analyzed the responses of these cells following the same procedures used for the original analyses in the corresponding experiments, except when otherwise noted (see below). Specifically, for Fig. 2f, top left and bottom left, we followed methods for Fig. 4d and e of Ref. 5, respectively; for Fig. 4b and c, we followed methods for Fig. 2a & b and Fig. 3a of Ref. 6, respectively. Since Refs. 5 & 6 did not analyze tuning curves conditioned on last walls, we used the average of the predicted tuning curves across all last walls. For Fig. 5d, top, middle and bottom rows, we followed methods for Fig. 1, 3rd row, 2A, and 3E of Ref. 7, respectively.

When the original analyses were not described in sufficient detail, we used the closest alternative. For detecting peaks in autocorrelograms for Fig. 4b, because the firing rates needed for thresholding were not available in Ref. 6, we used the same methods as Ref. 5. For computing the anisotropy of the autocorrelogram in the familiar environment, we then used the aspect ratio of the ellipse that best fits the six peaks closest to the center. Furthermore, because peak locations could only be estimated unreliably in unfamiliar environments, we computed the anisotropy there by multiplying the familiar environment’s anisotropy with the horizontal divided by the vertical scaling of the grid fields in the unfamiliar environment relative to the familiar environment shown in Fig. 4c. For ease of comparison with the rest of the results, we also computed the pooled responses for the rightmost pair of bars in Fig. 4c by first taking the proportion of the scaling of the grid fields relative to that of the environment within each category, *p*_s_. For example, scaling of 0.85, i.e., 15% compression, when the environment is scaled down 30% would yield *p*_s_ = 0.5, and scaling of 1.15, i.e., 15% expansion would yield *p*_s_ = −0.5. Then the relative proportion was converted back such that *p*_s_ *>* 0 would yield (1*/*0.7− 1) ·*p*_s_ + 1 and a negative one would yield (1 −0.7*/*1) ·*p*_s_ + 1, so that they can be compared to other bars. For consistency of simulations, we used a single set of grid modules across all simulated experiments to compute neural posteriors (see Predictions for grid cell responses). This means that the resulting grid fields did not necessarily match the scales of those measured in the particular corresponding experiments, which necessarily measured grid cell responses at only a limited subset of grid scales. Therefore, as our predicted grid fields only depended on a single optimized warping that was shared across all grid scales, for a fair comparison with experimental data, we generated grid cell responses using modules whose scales were specifically chosen such that they matched those reported in the original experiments. For this, using the optimized warping, we simulated seven grid cell modules whose scales formed a geometric series with a 1.42 quotient, with 100 grid cells each (with uniformly random phases). The smallest of these grid scales before warping was a free parameter Λ_0_. Note that experimentally observed grid scales correspond to the effective grid scales of our simulated grid cells after warping, as opposed to the ‘underlying grid scales’ that define the model (Λ_*i*_ in Eq. 28). Thus, we computed the effective grid scale of simulated grid cells in a familiar environment after warping, and only considered values for Λ_0_ that gave at least one rendered grid scale that was approximately within the experimentally observed range, defined as between 5 cm smaller than the minimum and 10 cm larger than the maximum reported in the corresponding experiment (Ref. 5: 24-55 cm, Ref. 6: 15-45 cm, and Ref. 7: 35-135 cm). For each experiment, among the values of Λ_0_ that satisfied this condition, we chose the one that gave the smallest discrepancy from the experimental results (see above). As the original recordings were from multiple animals, we chose the grid orientation of each simulated cell randomly with replacement from those in Ref. 5 (Extended Data Table 1, axis 1, in rows corresponding to the square environment: 36.4^°^, 11.3^°^, 58.0^°^, 63.4^°^, 17.2^°^, 14.9^°^, 17.5^°^, 23.2^°^, and 18.4^°^). Finally, we discarded simulated grid cells whose gridness fell below the minimum allowed gridness of the corresponding experiment (0.27 in the square environment^5^, 0.3 in the familiar environments^6^, or 0.4 in the familiar environment^57^).

#### Analysis of cue conflict in tethering experiments

Schematics in the bottom of Fig. 5a give intuitive explanation for main experimental measures (actual calculations used linear regression across 4 objects ×4 starting walls with the predicted shift appropriately changed depending on the starting wall^4^). The scaling of homing responses measures the ratio of the distance (A) between the two homing locations (orange) and the distance (B) between the target locations (or equivalent target locations, gray). The shift of responses measures the difference between the average of the homing locations (orange arrows) and the average of equivalent target locations (gray), with a sign that is defined to be positive when the shift is toward the starting wall.

## Extended Data Figures

**Extended Data Fig. 1.**
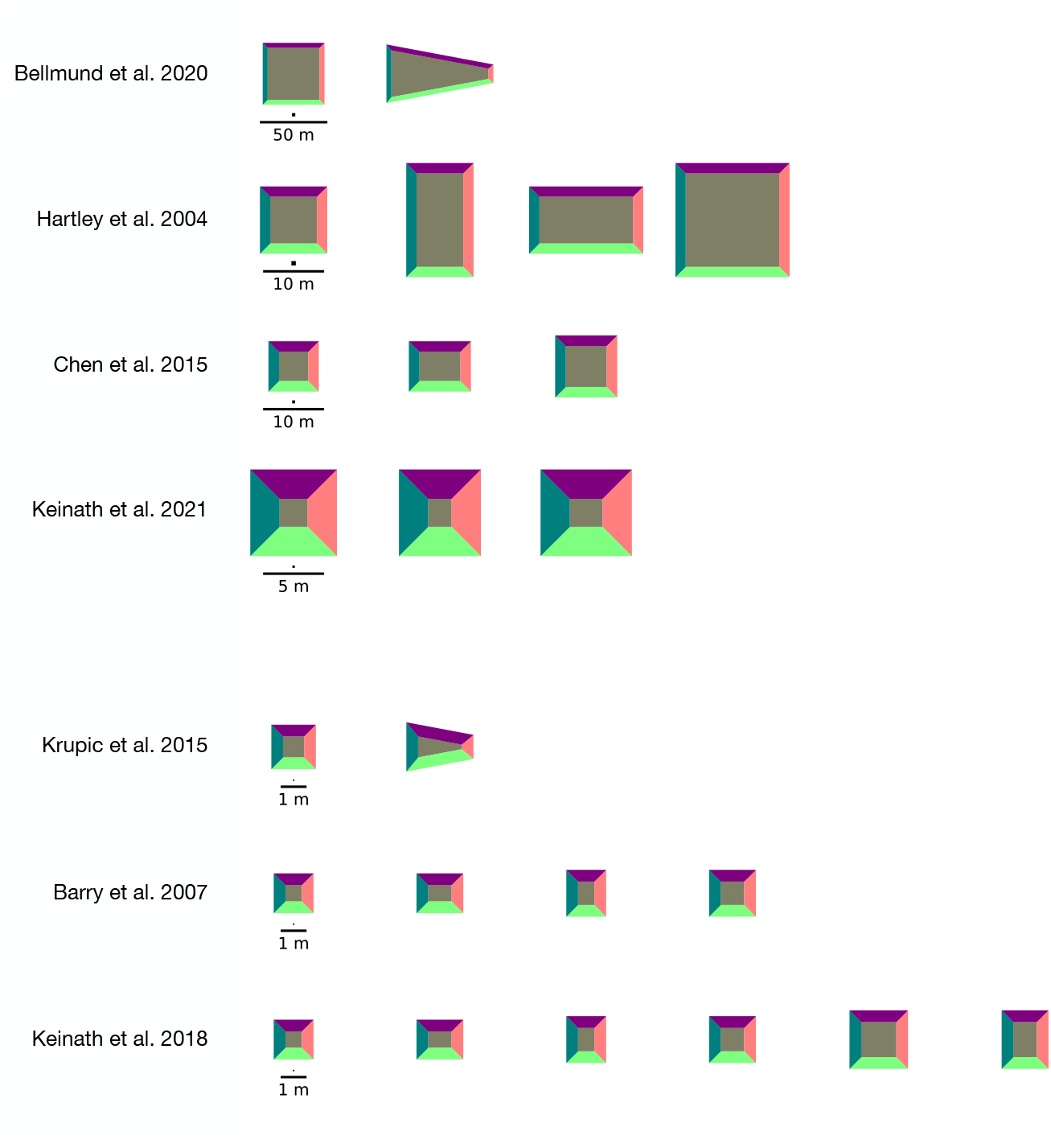
Simulated environments. Frame thickness around each environment shows wall height. The small square above the scale bar for each environment indicates δℓ, the spatial bin size used for the discretization of our simulations using that environment (see also Methods, Implementation details, and Table 2). Environmental geometries were reproduced from the original experiments (with humans, first four rows, or rats, last three rows): Bellmund et al. 2020^1^, used for results shown in Fig. 1c, Fig. 2d, Extended Data Fig. 2b; Hartley et al. 2004^2^, used for results shown in Fig. 3b-d, Extended Data Fig. 2c, Extended Data Fig. 7; Chen et al. 2015^3^, used for results shown in Fig. 3f, Extended Data Fig. 2d, Extended Data Fig. 8; Keinath et al. 2021^4^, used for results shown in Fig. 5b-c, Extended Data Fig. 2e (and also used as the basis for a more extremely expanded environment, for which results are shown in Extended Data Fig. 10); Krupic et al. 2015^5^, used for results shown in Fig. 1b, d-e, Fig. 2b, e-f, Extended Data Fig. 2f, Extended Data Fig. 3 left, Extended Data Fig. 5b, Extended Data Fig. 6; Barry et al. 2007^6^, used for results shown in Fig. 4, Extended Data Fig. 2g, Extended Data Fig. 3 middle, Extended Data Fig. 4, Extended Data Fig. 9; and Keinath et al. 2018^7^, in which the first four environments were from Barry et al. 2007^6^, and the last two were from Stensola et al. 2012^57^ (see caption of Extended Data Fig. 2 for an explanation). For Keinath et al. 2018^7^, the six environments together were used for results shown in Fig. 5d, Extended Data Fig. 2h, and Extended Data Fig. 11, and the last two were used for results shown in Extended Data Fig. 3 right.

**Extended Data Fig. 2.**
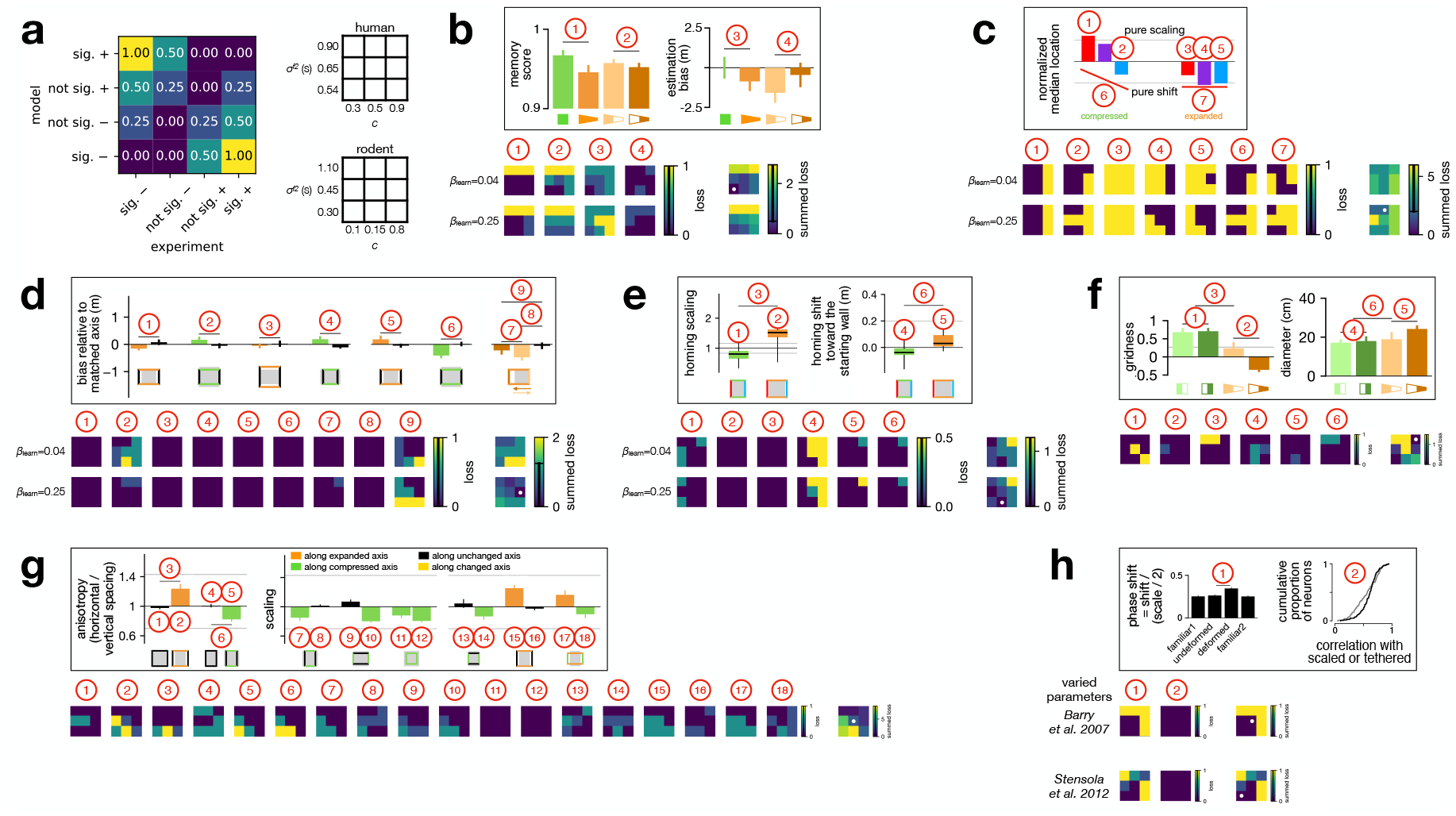
Experiment-specific parameters. **a**. Left: mismatch loss as a function of the sign and significance of the mismatch between modeled and experimentally measured statistical comparisons (see also Methods, Parameter choices). Right: 3*×*3 grids showing the values of two experiment-specific parameters (visual contrast *c*, and the relative variance of forward motion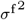) for which we report mismatch losses in the other panels (shown as smaller 3× 3 color maps), corresponding to human (top grid, and panels **b**-**e**) and rat experiments (bottom grid, and panels **f**-**h**). **b-h**. Values for experiment-specific ideal observer parameters were chosen by minimizing the mismatch loss for reproducing results in human homing (**b**-**e**) and rat grid cell (**f**-**h**) experiments. In each panel, inset at the top shows index to statistical comparisons of the experimental data as shown in the figure comparing them to BION results (see below). Heat maps in the bottom show mismatch loss for each statistical comparison as a function of experiment-specific parameters (*c*, 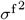 and also memory fidelity *β*_learn_ for human experiments, see Table 2 and Methods for further details). Last heat map in each panel shows summed mismatch loss across all comparisons in that panel, with white dot indicating the parameter combination chosen for simulating each experiment (associated with minimal summed mismatch loss). When the summed loss did not have a unique optimum, we chose the parameter combination with which BION produced results that were visually most similar to the corresponding experimental results. **b**. Simulations for Bellmund et al. 2020^1^, results shown in Fig. 2d. **c**. Simulations for Hartley et al. 2004^2^, results shown in Fig. 3b-d, Extended Data Fig. 7, inset shows Fig. 3d. **d**. Simulations for Chen et al. 2015^3^, results shown in Fig. 3f, Extended Data Fig. 8, inset shows Extended Data Fig. 8a-c. **e**. Simulations for Keinath et al. 2021^4^, results shown in Fig. 5b-c (chosen parameters also used for Extended Data Fig. 10), inset shows Fig. 5b. **f**. Simulations for Krupic et al. 2015^5^, results shown in Fig. 2e-f, Extended Data Fig. 3 left, Extended Data Fig. 5b, Extended Data Fig. 6, inset shows Fig. 2f. **g**. Simulations for Barry et al. 2007^6^, results shown in Fig. 4, Extended Data Fig. 3 middle, Extended Data Fig. 4, Extended Data Fig. 9, inset shows Fig. 4b-c. **h**. Simulations for Keinath et al. 2018^7^, results shown in Fig. 5d, Extended Data Fig. 3 middle and right, Extended Data Fig. 11, inset shows Fig. 5d. Note that, in most cases, each experiment contributed data to one experimental paper whose results we compare with the predictions of BION (and report in specific figures of the main paper, corresponding to different panels here, see above). An exception to this is the experiment of Barry et al. 2007^6^, whose data were analyzed both in the same paper (see our Fig. 4 and Extended Data Figs. 3, 4 and 9, and panel **g** here) and by Keinath et al. 2018^7^ (see our Fig. 5 and Extended Data Figs. 3 and 11, and panel **h** here). Thus, we chose a single set of parameters for the Barry et al. 2007^6^ experiment, which then contributed to our comparisons of BION to the the results reported by both Barry et al. 2007^6^ and Keinath et al. 2018^7^ (and the corresponding losses shown in panels **g** and **h** shown here). Conversely, Keinath et al. 2018^7^ reported the combined analyses of two data sets: one collected by Barry et al. 2007^6^, and another one collected by Stensola et al. 2012^57^. Thus, we allowed different parameter choices for these two experiments, and show the dependence of the comparison of BION to the results reported by Keinath et al. 2018^7^ (and the corresponding loss in panel **h**) as a function of both sets of parameters. As Fig. 1b-e and Fig. 2b illustrate BION’s behavior, rather than compare it to experimental data, there are no losses shown for them. The parameters used for these illustrative simulations were: *c* = 0.2, 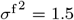 s for Fig. 1b,d-e, *c* = 0.3, 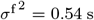, *β*_learn_ = 0.1 for Fig. 1c, and *c* = 0.2, 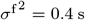 for Fig. 2b.

**Extended Data Fig. 3.**
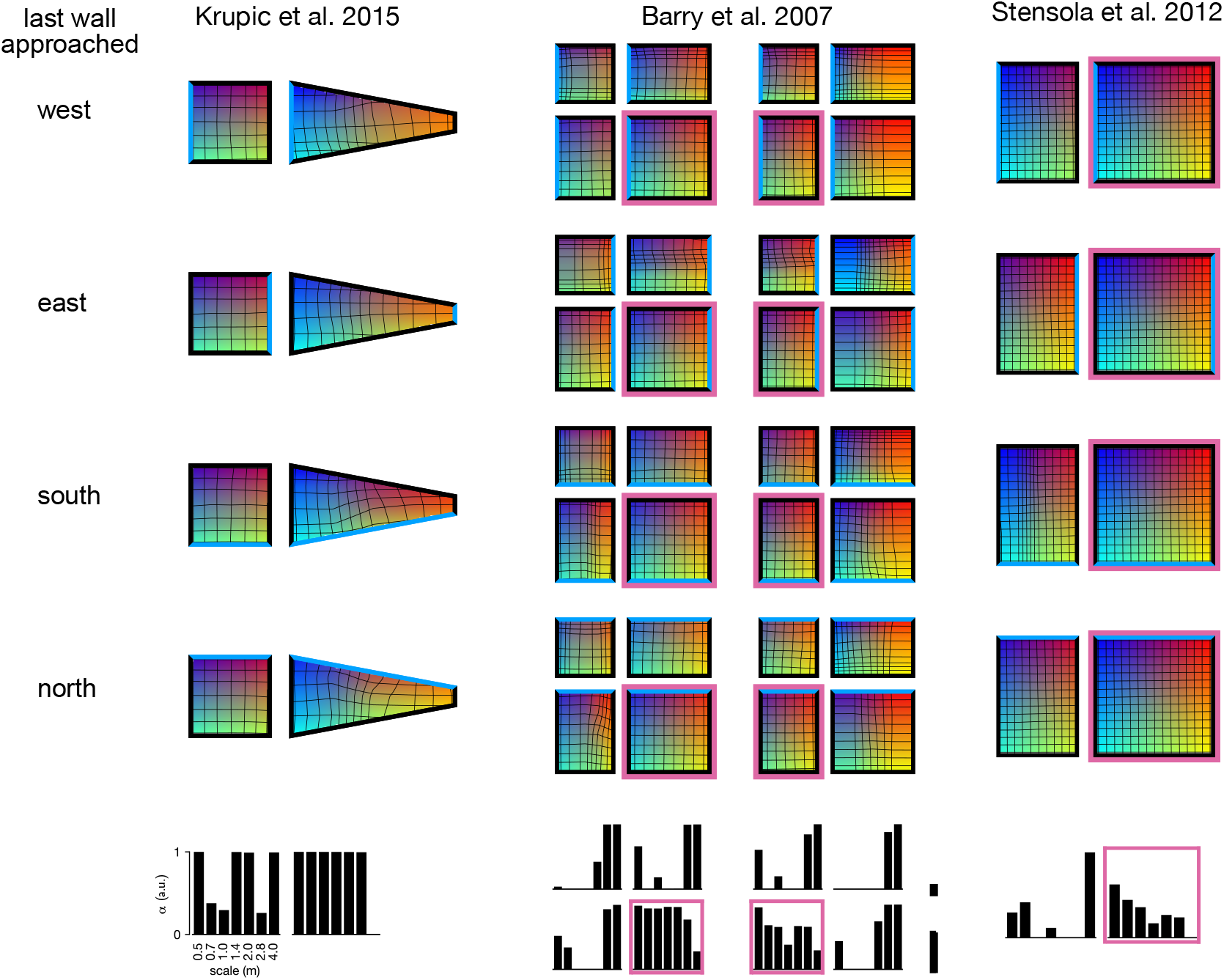
Grid cell parameters optimal for matching neural to ideal observer posteriors. Top four rows: optimized warpings (shared across all cells). Isocontours of the x and y coordinates in the believed environment (**h** ℓ; 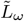 in Eq. 28) that represent how optimized encoding grid fields (in the decoding-based approach) are warped. The colors indicate the x (blue to red, from left to right) and y (black to green, top to bottom) values. Each row represents the isocontour when the agent approached each wall last (from top to bottom: west, east, south, or north; marked with blue). Supercolumns from left to right represent warpings used for simulating Krupic et al. 2015^5^, Barry et al. 2007^6^, and Stensola et al. 2012^57^, respectively. (See caption of Extended Data Fig. 2 for an explanation of how the Barry et al. 2007^6^ and Stensola et al. 2012^57^ experiments relate to the results reported by Barry et al. 2007^6^ and Keinath et al. 2018^7^.) Pink outlines indicate familiar environments in experiments where unfamiliar environments were used. Bottom row: optimized *α* values for each grid cell module (x-axis shows corresponding grid scales before warping; Eq. 31).

**Extended Data Fig. 4.**
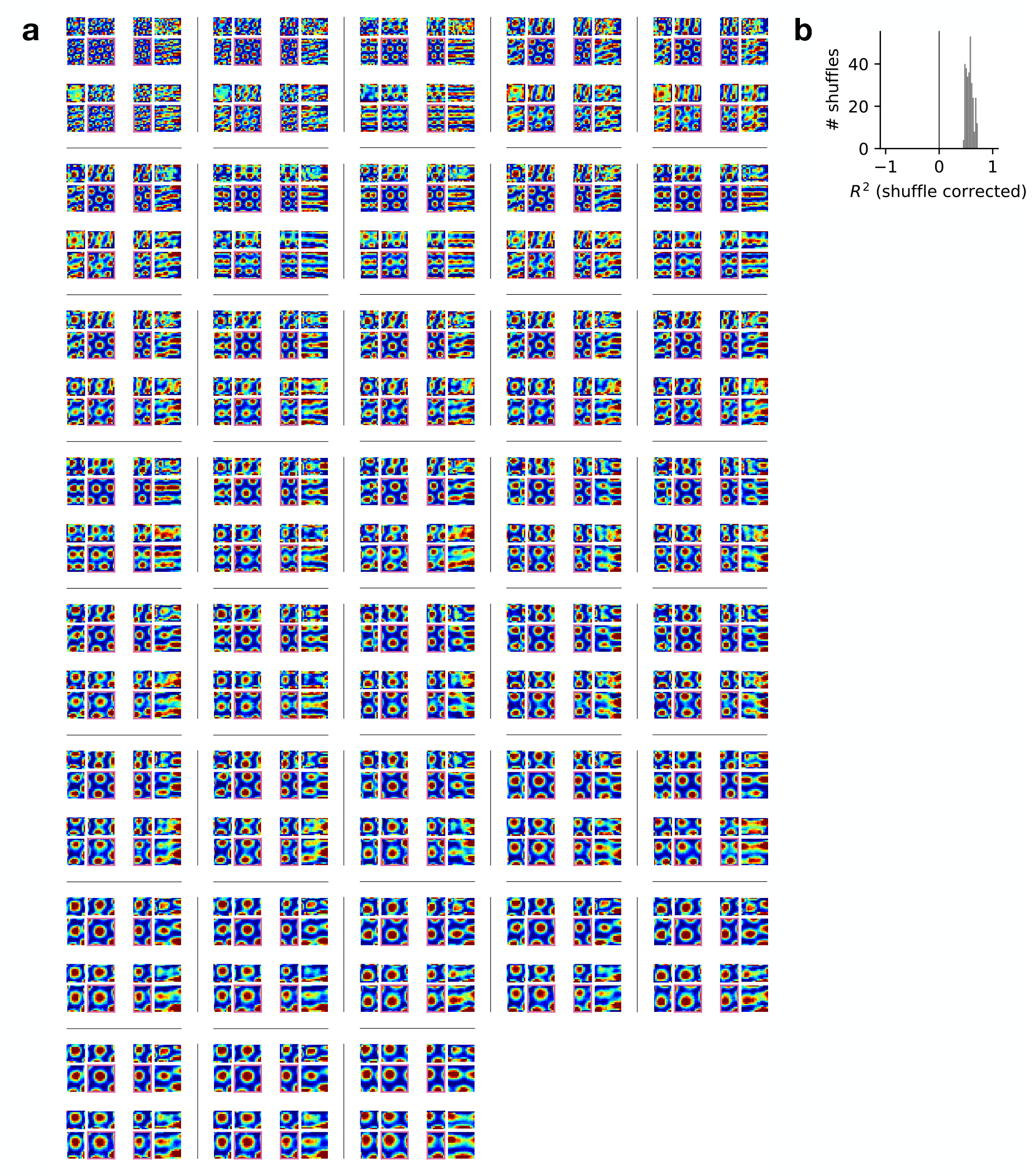
Encoder-decoder duality. **a**. Comparing tuning curves between the decoding- and encoding-based approaches. Each 2 ×2 panel (between black lines) shows the tuning curves of the same cell: decoding-(top row) vs. encoding-based tuning curves (bottom row), when the large square environment (left columns) or the vertical rectangular environment (right column) is familiar (pink outlines, as in Fig. 4 and Extended Data Fig. 9). Cells are ordered from small to large grid scale from left to right, and top to bottom (as in Extended Data Fig. 9). Color scales from the 10th and 90th percentiles of firing rates within each panel. **b**. Match between encoding-based and decoding-based tuning curves (Eqs. 27 and 28), measured as variance explained (*R*^2^). To control for spurious correlations, the *R*^2^ obtained by optimizing the encoder to match time-shuffled decoding-based firing rates is subtracted from the *R*^2^ obtained by optimizing the match with the original (non-shuffled) decoding-based firing rates is shown (mean*±*SD 0.57*±*0.06).

**Extended Data Fig. 5.**
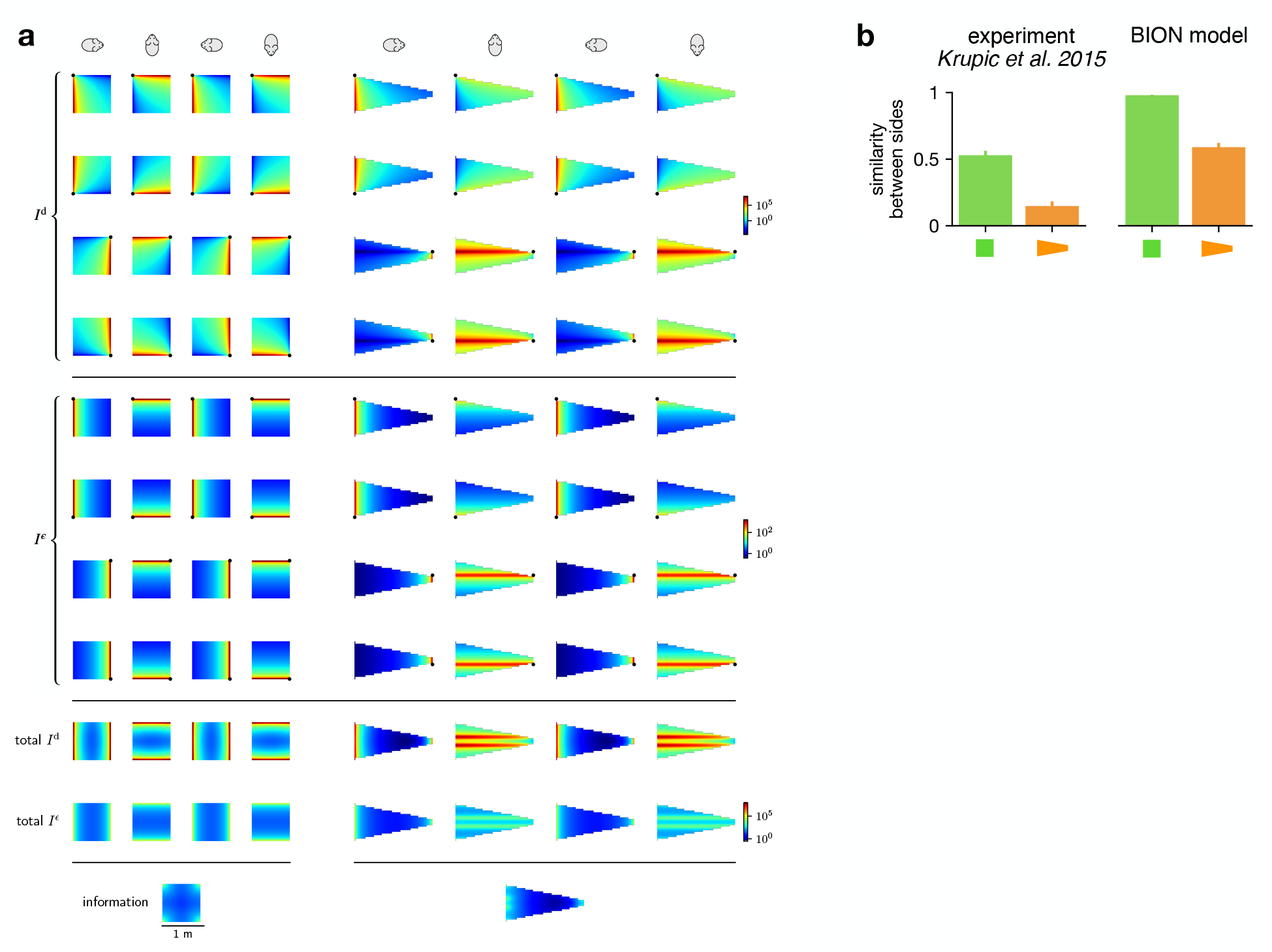
Visual information and grid field similarity in square and trapezoid environments. **a**. Parameter-free analytical prediction of visual information in square (left) and trapezoid (right) environments (such as those used by Krupic et al. 2015^5^ and Bellmund et al. 2020^1^, though scales differ). For simplicity, here we assume point-like landmarks in the corners, instead of walls, and 360^°^-vision (with heading direction only relevant for determining the orientation of the imaging plane, and thus for determining which allocentric direction is along or orthogonal to the egocentric line of sight), explaining why information maps across the odd columns and across the even columns of each environment are identical. *Rows 1–4*. Information about location along line of sight for a given heading *θ* (columns) provided by each corner of the environment (rows: *c* = north-west/south-west/north-east/south-east, marked with a black dot), computed using Eqs. 1 and S19: 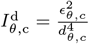. *Rows 5–8*. Same as rows 1–4, but for information about location along the direction that is orthogonal to line of sight, computed using Eq. S22: 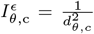. Note that we have ignored constants of proportionality in Eqs. 1, S19 and S22 as they are identical for 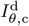 and 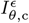, and as such would simply scale all the results shown in this figure. *Rows 9 & 10*. Information along (row 9) and orthogonal to (row 10) line-of-sight for a given heading (columns) with information integrated across four corners: total 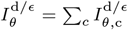. Note that we sum *information* across all four corners because we assume that each would have been visible sufficiently recently. *Row 11*. Information that corresponds to the uncertainty averaged across headings: 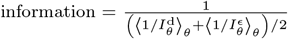. The average uncertainty across heading directions expresses the expected Euclidean error in location estimation at a given location, assuming that the agent is equally likely to have any of the heading directions. Note that this is consistent with the way spatial information maps in Fig. 2b were computed (see also Methods, Variation of visual information within an environment). Indeed, the results we show here with this parameter-free analytical prediction are qualitatively similar to those produced by numerical simulations of the full BION model, shown in Fig. 2b: total information is higher in the square than in the trapezoid environment, and overall higher in the broad than in the narrow side of the trapezoid. **b**. Similarity of grid fields between the two halves of each environment in data^5^ (left) and BION (right).

**Extended Data Fig. 6.**
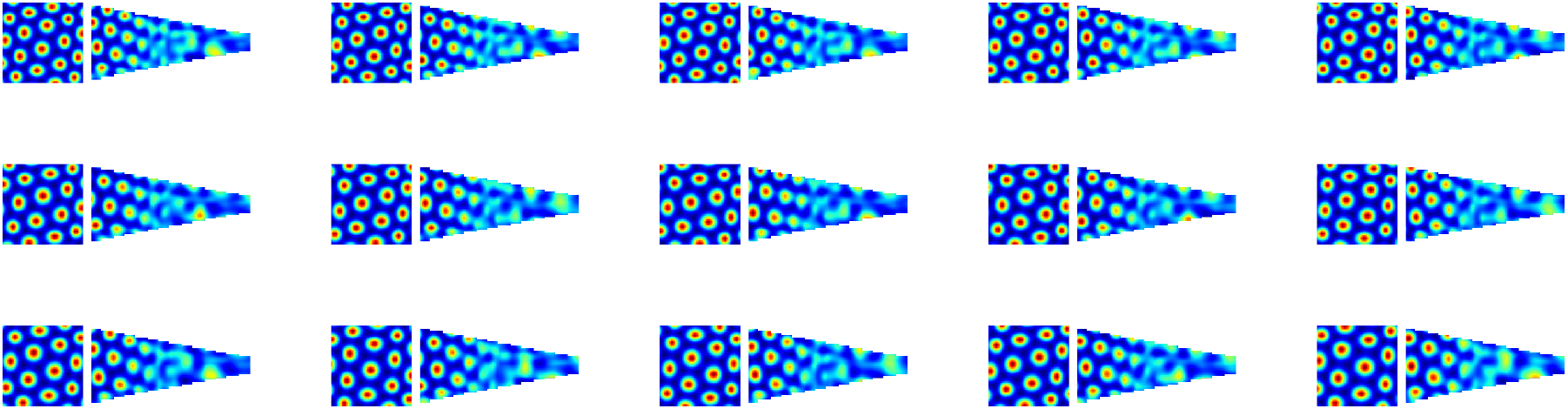
All grid fields analyzed from the model fit to Krupic et al. 2015^5^ (extending Fig. 2e).

**Extended Data Fig. 7.**
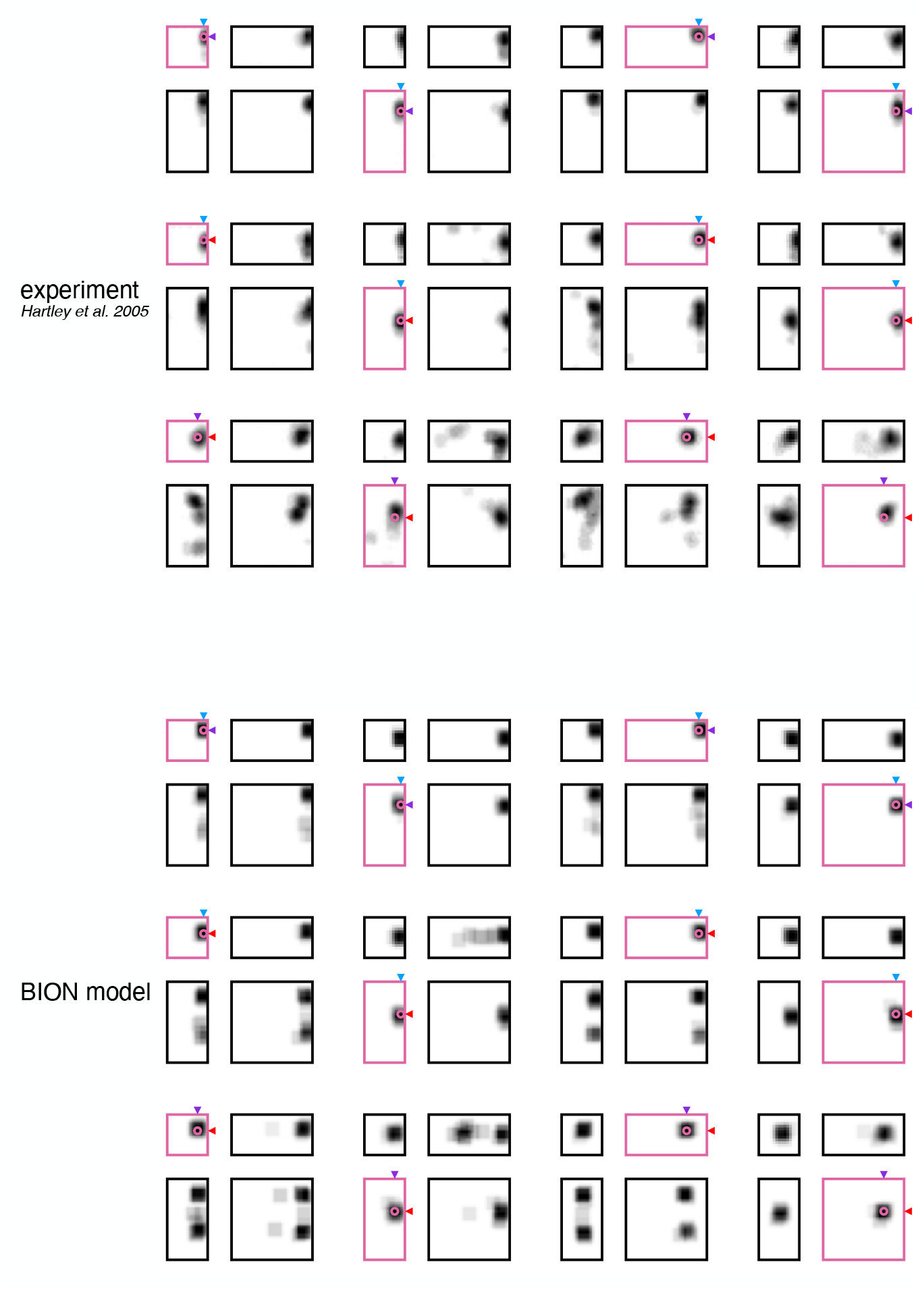
Full results for Hartley et al. 2004^2^ (extending Fig. 3a). Blue, purple, and red arrowheads show when the target’s coordinate along the corresponding axis is near, intermediate, and far from the nearest wall along that axis, respectively. Pink outlines indicate familiar environments.

**Extended Data Fig. 8.**
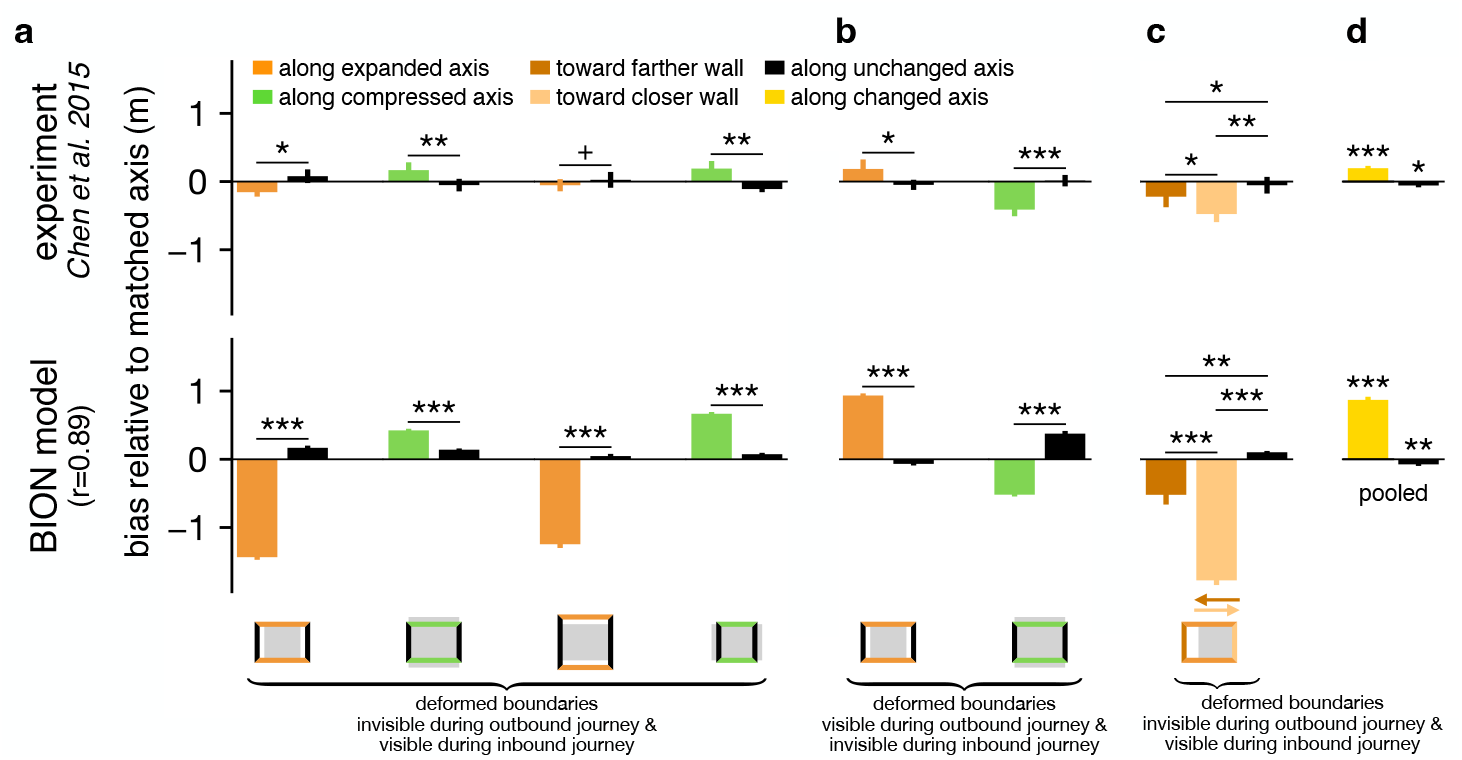
Full results for Chen et al. 2015^3^ (extending Fig. 3f). **a**. Experiments 1 & 2 as in Fig. 3f. Deformed boundaries are invisible during outbound journey & visible during inbound journey. **b**. Experiment 3. Deformed boundaries are visible during outbound journey & invisible during inbound journey. **c**. Experiment 4. Deformed boundaries invisible during outbound journey & visible during inbound journey as in **a**., except that only the farther wall moves away and the closer wall stays at the same location. **d**. Results pooled across **a**–**c** (also shown in Fig. 3f).

**Extended Data Fig. 9.**
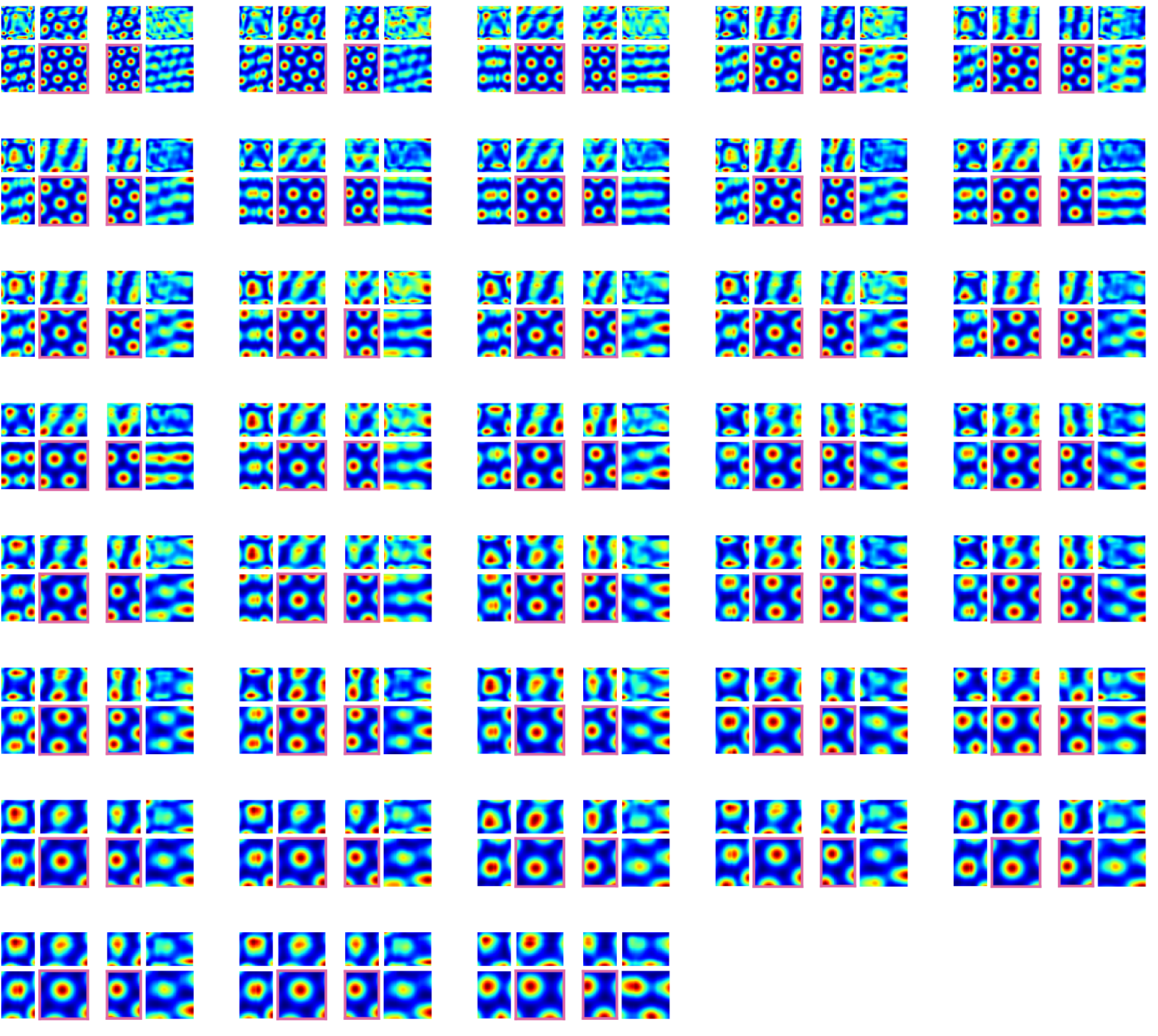
All grid fields analyzed from the model fit to Barry et al. 2007^6^ (extending Fig. 4a). Pink outlines indicate familiar environments.

**Extended Data Fig. 10.**
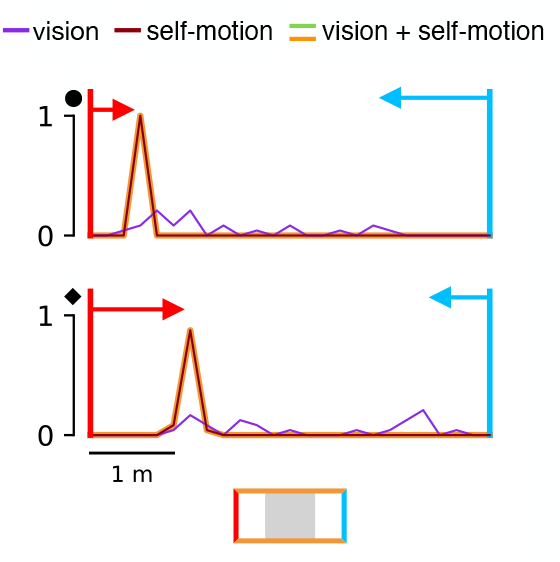
Predictions for tethering (extending Fig. 5c). Prediction of results under more extreme expansion (1.58x, rather than 1.17x; see Fig. 5c) for Keinath et al. 2021^4^.

**Extended Data Fig. 11.**
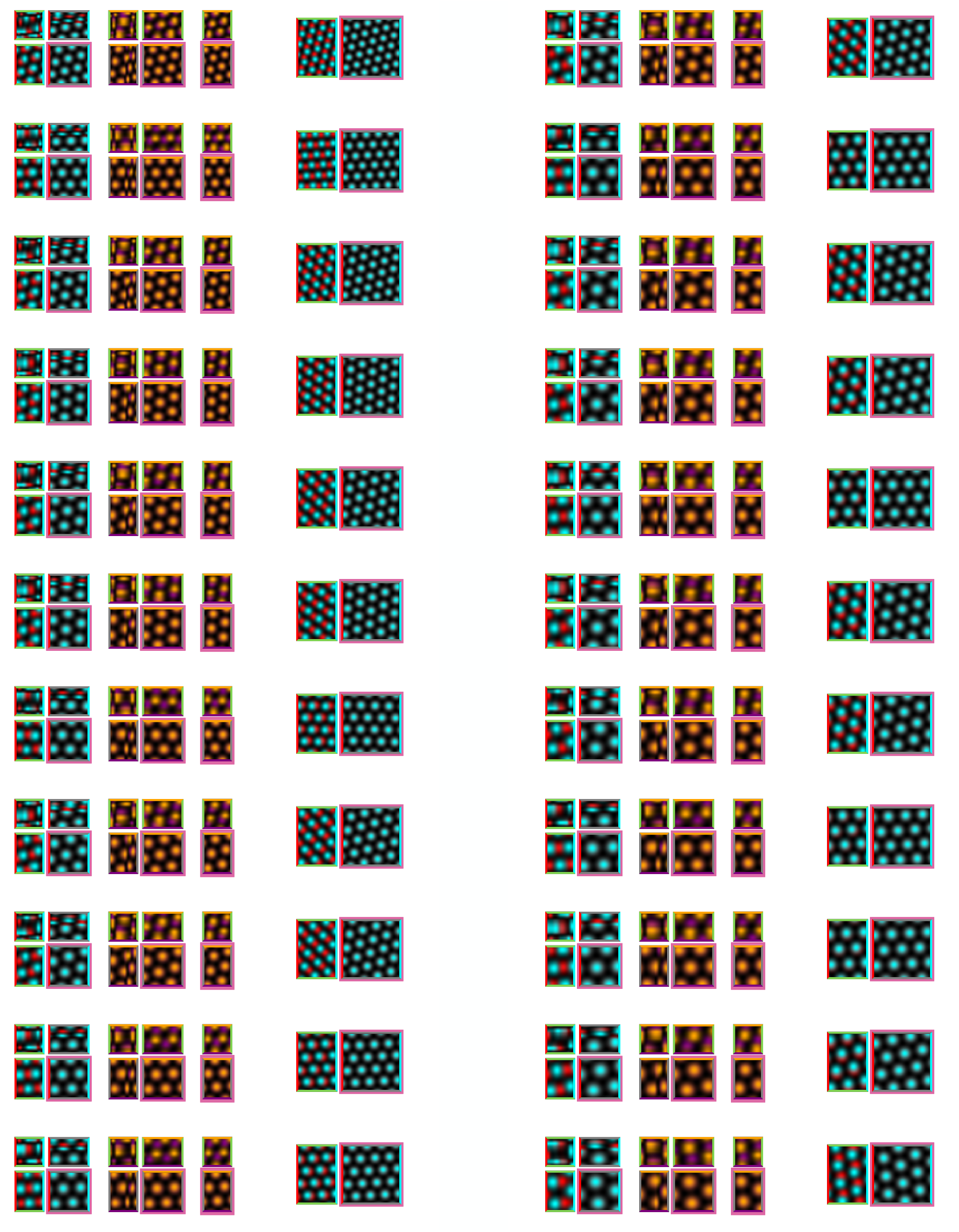
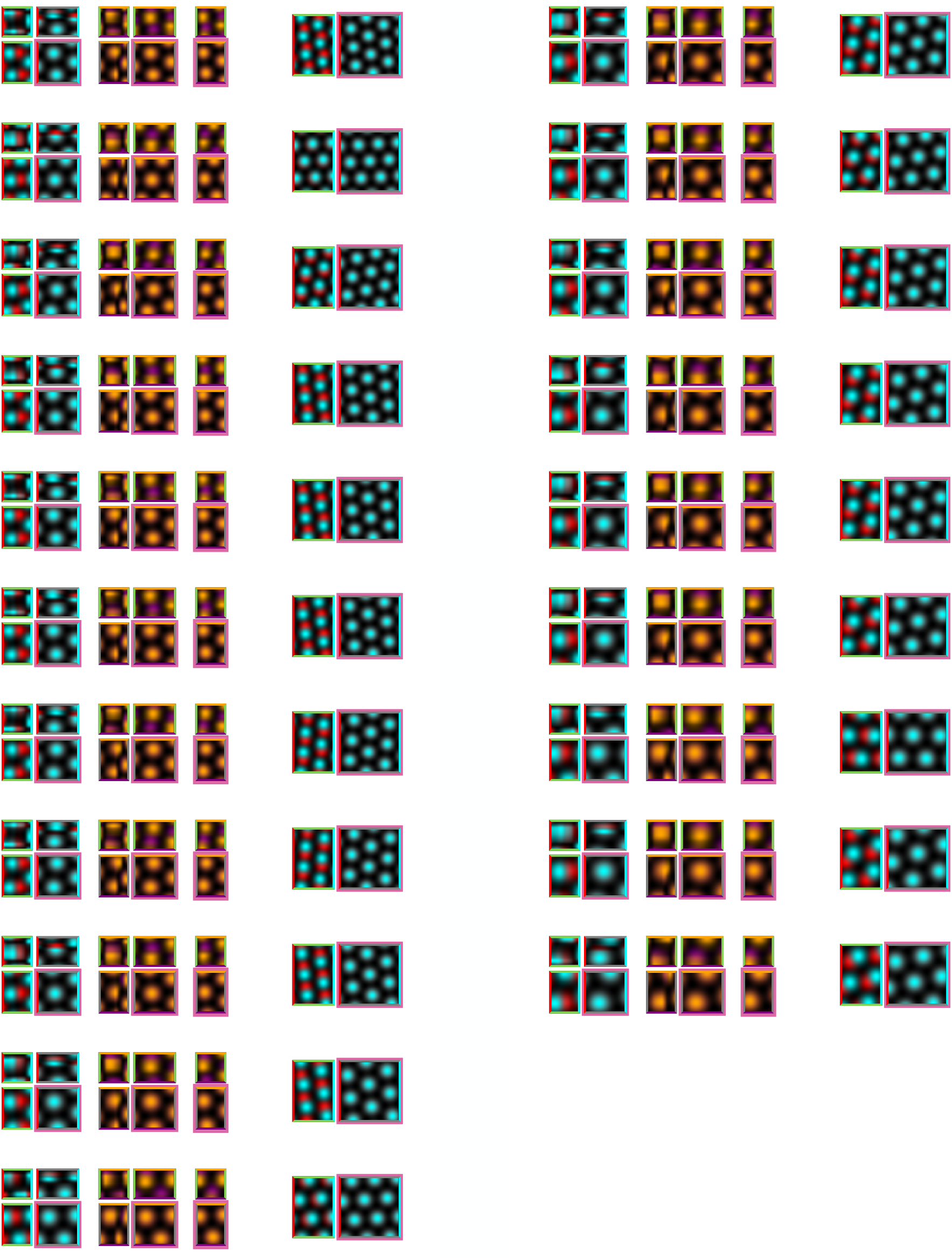
All grid fields analyzed from the model fit to Keinath et al. 2018^7^ (extending Fig. 5d). Pink outlines indicate familiar environments.

## Supplementary Material

### SM1. Contribution of binocular disparity to visual information

In our main analyses, we use cyclopean vision and ignore binocular disparity. Here we show that the contribution of binocular disparity to visual information about depth is small in the experiments we considered. Based on Eq. 1 (see also Eqs. S19 and S29), we derive the information afforded by binocular disparity when the interpupillary distance is 2Δ as:

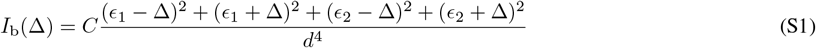

where *C* is a constant, *ϵ*_1_ and *ϵ*_2_ are the eccentricities (from the midline of the agent looking ahead) of the two corners of a wall that is in front of the agent and orthogonal to the line of sight. The four terms in the numerator correspond to the effective eccentricities of the two corners of the wall from the line of sight of either of the two eyes. (Note that this is actually an upper bound on binocular information, and thus a worst-case scenario for our analysis here, obtained when the information provided by the retinal eccentricity of either corner, and especially in either eye, is independent and so those individual information terms simply sum linearly.) We use *W* to indicate the width of the wall, such that *ϵ*_2_ = *W− ϵ*_1_ and 0 ≤ *ϵ*_1_, *ϵ*_2_ ≤ *W*. Likewise, we derive the information afforded without binocular disparity as the special case of the above when Δ = 0:

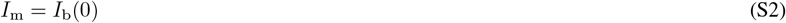

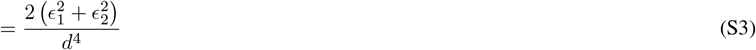

Note that we have kept the factor 2 in the numerator, because as long as the information without binocular disparity is just a constant factor less than that with binocular disparity, this difference can always be offset by reducing the overall amount of effective noise in the retina. The main concern is whether the scaling of information with environmental geometry differs in the two cases. For this, we define the relative gain in information from binocularity *b* as:

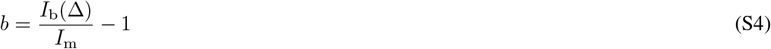

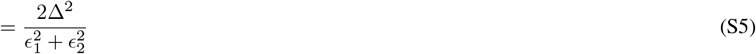

It is easy to see that, as a function of *ϵ*_1_, *b* has a maximum at 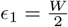:

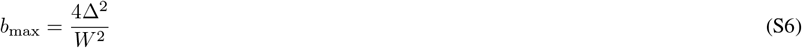

In human experiments, Δ ≈ 0.04 m^62^ (conservatively large estimate, such that the estimated binocular advantage is maximized) and the smallest width of a wall considered is *W* = 2 m (compressed room of Ref. 4), giving *b*_max_ = 0.0016, i.e. less than 0.2% of advantage. In rat experiments, Δ ≈ 0.012 m^63^ (using the width of the rat skull, specifically the bizygomatic width, as the upper bound for their interpupillary distance, again, as a conservatively large estimate) and the smallest width of a wall considered is *W* = 0.2 m (narrow side of Ref. 5), giving *b*_max_ = 0.0144, i.e. less than 1.5% of advantage. These differences (*<* 1.5%) are almost an order of magnitude smaller than the difference in information between locations within an environment or between environments that we obtained (*>* 10%, Fig. 2b). Thus, we conclude that considering binocular vision would not change our results qualitatively.

### SM2. Encoder-decoder duality

In order for BION to make predictions about neural responses, we need to define a mapping between the ideal observer’s posterior (the primary prediction of BION) and neural responses. In the main text, we defined this mapping indirectly via a *decoding-based approach*: we started by defining a transformation from neural responses to probability distributions (this is what we called a ‘decoder’), and then made neural predictions by finding the tuning curves (under the constraints of our warped grid field parametrization) that generated neural responses for which the decoded probability distributions (the neural posteriors) best matched the ideal observer’s posteriors (Eqs. 27-33 and 35). However, this approach relied on defining tuning curves, and thus neural responses at any time, as functions of the true state of the animal, 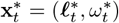 (where 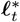 is the true location of the animal at time *t*, and *ω*_*t*_^∗^ is the identity of the last wall it approached). This may seem paradoxical as it seemingly requires grid cells to have access to the true state of the animal, while the fundamental tenet of our model is that the animal itself does not have access to its true state and can only infer a posterior distribution over it. Moreover, in this approach, for scaling experiments in which there is a mismatch between the new, scaled environment in which the animal actually is, and the familiar environment in which the animal believes itself to be, we had to re-optimize the tuning curves that generate neural responses (the ‘encoding’ tuning curves) in a way that they were defined in the new environment (so that they could be compared to empirically measured tuning curves which are necessarily defined in the new environment). This makes it appear as if an instantaneous optimization process needed to take place the moment the animal entered the new environment, and grid cells not only needed to have access to the animal’s true state but also to the true environment in which it is, while the ideal observer analysis assumes that the animal believes it is still in the familiar environment. Thus, the question is: how can grid cell firing appear to depend on information (about true state and true environment) to which the animal (and thus our ideal observer model) has no access, and how can it be re-optimized on entry to a yet-unknown environment?

To resolve this apparent paradox, we show that the decoding-based approach of the main text is equivalent to an *encoding-based approach* that determines neural responses directly as a function(al) of the ideal observer’s posterior over locations, without any reference to the true state or environment of the animal, and without any re-optimization taking place when the animal unknowingly enters a new environment.

To develop mathematical intuition for this duality, recall from Eq. 31 that we decode the neural posterior from neural responses as follows (simplifying notation to reduce clutter):

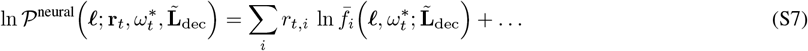

where ℓ is (a candidate) decoded position (in the believed familiar environment), **r**_*t*_ is the vector of grid cell firing rates at time *t* (generated based on the encoding tuning curves, which are functions of ℓ^∗^ and *ω*^∗^ in the actual environment), and 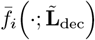 are the (normalized) decoding tuning curves (defined in the believed environment) – and we made their dependence on both ℓ and *ω* explicit. Once neural responses (i.e. the gain and warping of encoding tuning curves that give rise to them, and to which decoding tuning curves are tied in familiar but not in novel environments) have been optimized (for the objective of Eq. 33), neural posteriors ought to match the ideal observer’s posterior at least approximately (again after simplifying notation):

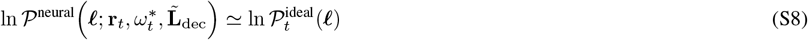

Substituting this to Eq. S7, and rewriting the resulting equation slightly in order to get rid of the unwieldy (log) normalization factor in Eq. S7@, gives

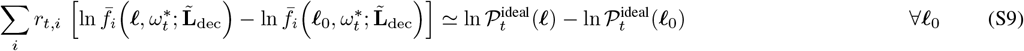

In order to take an encoding-based approach, we need to express neural responses as some function of the ideal observer’s posterior from this equation. For this, we rewrite Eq. S9 in discretized space as

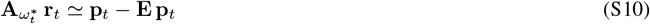

where

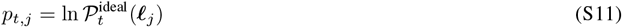

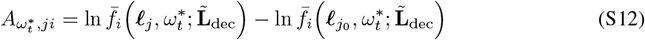

and 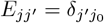, with *j*_0_ being the index of the discretized location bin corresponding to ℓ_0_, and **p**_*t*_. For simplicity, we assume that the number of location bins in discretized space is larger than the number of neurons. (This can always be assumed, as in principle we require the ideal observer’s and neural posteriors to match at an arbitrarily fine spatial scale.) This allows us to solve Eq. S10 for neural responses as

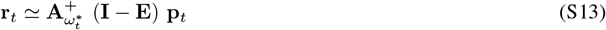

where

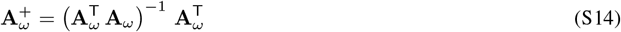

is the left pseudoinverse of **A**_*ω*_. Therefore, Eq. S13 suggests that neural responses that are near-equivalent to those obtained by our original decoding-based approach could also be obtained as a direct function(al) of the ideal observer’s posterior, without any reference to the true location (or environment) of the animal (albeit formally still requiring knowledge of 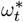). In particular, Eq. S13 suggests that such neural responses can be obtained as a simple linear function of the logarithm of the ideal observer’s posterior.

To illustrate the encoder-decoder duality, we revisit the rescaling experiment of Ref. 6, and – motivated by Eq. S13 – for each simulated grid cell, we apply a fixed linear function (the encoder) to the logarithm of the ideal observer’s momentary posterior over locations (always defined in the familiar environment) to compute the cell’s momentary firing rate (in the familiar, or in one of the scaled environments). As Eq. S8 (and thus Eqs. S10 and S13) are only approximate, and because we require an encoder that is independent of 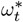, the exact coefficients for this linear encoding function may not be as simple as those suggested by Eqs. S12 and S14. Thus, we obtain the encoding coefficients by linear regression of the (discretized) log posterior against the neural responses predicted by our original decoding-based approach (Fig. 4 and Extended Data Fig. 9). Importantly, we use a single set of 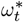-independent encoding coefficients across all four environments (familiar and three scaled environments), so no re-optimization takes place when the animal unknowingly enters a new environment. With these coefficients, tuning curves constructed from the responses generated by the encoding-based approach are near-identical to those predicted by our original decoding-based approach (Extended Data Fig. 4).

In summary, the encoding-based approach described here allows us to define a transformation from (the history of) sensory inputs to neural responses in two steps: in the first step (the ideal observer), we transform sensory inputs into posteriors, and in the second step (the encoder), we transform posteriors into neural responses. Of course, neither the decoding-, nor the encoding-based approaches provide a mechanistic account of how neural circuits and plasticity mechanisms actually achieve this overall transformation from sensory inputs to neural responses. (For examples of how mechanistic models performing such probabilistic computations can be constructed, see e.g. Refs. 41 and 64.) Nevertheless, the match between the encoding- and decoding-based tuning curves we find means that the particular transformation we obtain has the appealing property that a simple linear decoder that is based on the empirical tuning curves of grid cells in the familiar environment is able to decode neural responses into (approximately) correct (log-) posteriors (in the believed familiar environment) no matter what the true environment is. Over time, with sufficient experience, information about empirical tuning curves in the familiar environment can be accumulated in downstream areas even without access to the true state of the animal at any one time^65^, thus providing the necessary information for constructing such a decoder. In this sense (the parametric form of the decoder is simple linear, and its parameters can be empirically estimated with experience), the encoder-decoder duality implies that grid cell responses (approximately) encode the ideal observer’s posterior such that the resulting code is easily decodable.

### SM3. Visual uncertainty and environmental geometry

#### Information about the distance of a point landmark in retinal eccentricity

While all the results in the main text are obtained from numerical simulation using the full image-computable ideal observer model, we also derived a simple analytic relationship between Fisher information about location and environmental geometry to gain analytic insights, and confirmed its correspondence with the numerical simulations (Fig. 2a). Here we start by deriving the Fisher information 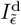 that the retinal (pinhole) projection of a single point landmark provides about its distance from (the pinhole aperture of) the observer *along* the line of sight, *d*, when it is at eccentricity *ϵ* (distance *from* the line of sight; see also Fig. 2a). Note that if the map of the environment, i.e. the allocentric location of the landmark, is known – as we assume to be the case – then this is the same information that this landmark provides about the observer’s allocentric location along its line of sight.

Without loss of generality, we assume that the distance of the imaging plane from the pinhole aperture is 1 (so that landmark distances are effectively measured in units of imaging plane distance). Thus, due to the basic geometry of a pinhole projection, this landmark would be seen at a retinal eccentricity (i.e. in the imaging plane, which we assume here to be infinitely large for simplicity, so that we do not need to treat the special case when the pinhole projection falls off the imaging plane) that is

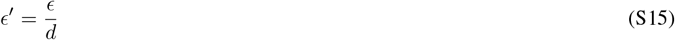

However, we also assume Gaussian blur of variance *σ*^2^ in the imaging plane, so that the eccentricity at which the observer actually observes the landmark is

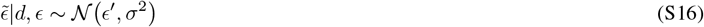

(Note that retinal eccentricity is the only source of information about the distance from a point landmark for a pinhole projection.) The Fisher information is then given as

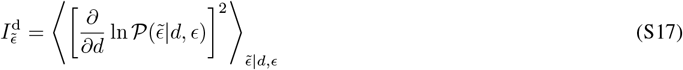

Because the relationship between *d, ϵ*, and *ϵ*^*′*^ is deterministic (Eq. S15), and the observed variable only depends on *ϵ*^*′*^ directly (Eq. S16), this can be rewritten as:

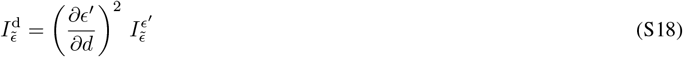

where 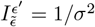 is the Fisher information in 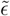 about *ϵ*^*′*^ (based on Eq. S16), which yields

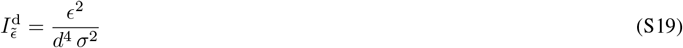

which is what we show (up to the constant factor of proportionality 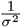) as Eq. 1 in the main text. This equation expresses the simple intuition that given a finite-resolution retina, the information about the distance to a landmark depends on how far its image moves on the retina per unit translation along the line of sight. For example, if the eccentricity is zero then the image does not move on the retina as the observer approaches the landmark (no information about distance). More generally, the closer the landmark is, and the larger its eccentricity, the more its retinal image moves for a unit change in the observer’s distance to it.

Similarly, the information about the true eccentricity of the landmark (or, vice versa, the distance of the observer from the landmark in the direction orthogonal to its line of sight) is given as

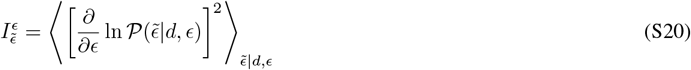

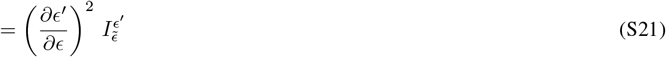

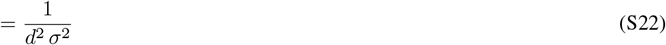

For completeness, it is useful to also compute the off-diagonal element of the Fisher information matrix, which is

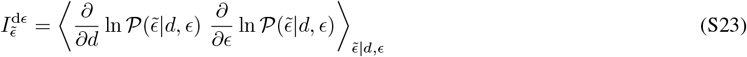

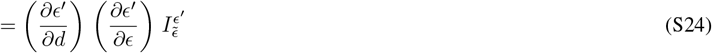

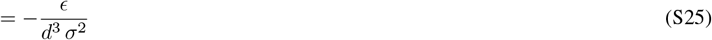

#### Information about the distance of a point landmark in the response of a retinal eccentricity-tuned neural population

While in the previous (sub)section we assumed that retinal eccentricity as such was directly observable, we now treat the more realistic scenario when instead only the pattern of activation across an array of retinal receptors is observable. We assume a standard population coding model, according to which each neuron *i* has a tuning curve for the position of a point stimulus (the projection of the point landmark) in the retinal plane. Retinal position is expressed in radial coordinates (a combination of retinal angle and eccentricity), and the tuning curve is a product of identical Gaussians for each radial coordinate, so e.g. for *ϵ*^*′*^, it is:

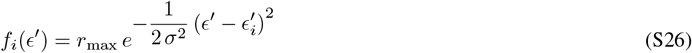

where 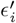 is the preferred eccentricity of the neuron, and *r*_max_ and *σ*^2^ are respectively its maximal firing rate and the (half)width of its tuning curve (the same for all neurons, hence the absence of the index *i*). (Note that even though the angle component of position is a circular variable, we assume that *σ* in that space is sufficiently small that the tuning curve can still be characterized with a normal, i.e. non-circular, Gaussian.) We assume independent Poisson noise across neurons, and a dense array of neurons that cover all two-dimensional preferred retinal positions uniformly with density *ρ* (note that a uniform coverage along the angle dimension requires that the number of neurons with a given preferred eccentricity scales with that preferred eccentricity). The Fisher information that the neural response of such a population provides about *ϵ*^*′*^ during a time window of *T* is given as^58^:

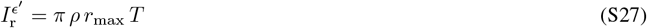

(Note that, as explained in Ref. 58, 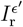 happens to be independent of the width of the tuning curve, *σ*, in the special case we are considering here, i.e. that of a neural population tuned to two stimulus features with a uniform tuning density. This independence is not essential for the following, so our final result would remain the same, up to a constant factor that may depend on *σ*, even if neurons were tuned to more or less stimulus dimensions.) Finally, as in Eqs. S18 and S19, we can obtain the Fisher information about *d* simply as

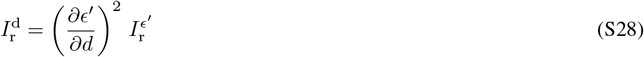

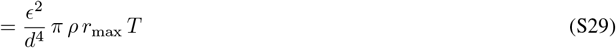

which is also equivalent to Eq. 1 in the main text, up to a constant factor of proportionality (which in this case is 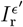). Similarly, the Fisher information about *ϵ* is simply

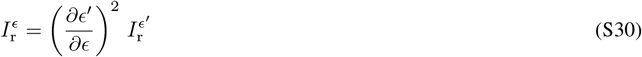

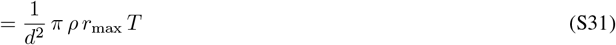

and the off-diagonal element of the Fisher information matrix is

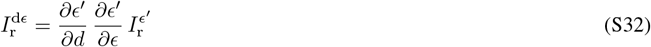

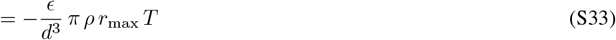

#### Interpretation of Fisher information

Note that the equations above (Eqs. S19, S22, S25, S29, S31 and S33) express the Fisher information available in a snapshot of visual information when only a single point-like landmark is available. Fisher information is relevant because its inverse expresses (an upper bound on) uncertainty (expressed as posterior variance for a uniform prior). However, there is a seemingly paradoxical aspect of the equations we derived above: although they show that Fisher information (each element of the Fisher information matrix) is finite as such, the uncertainty that this Fisher information matrix implies (the posterior covariance) actually becomes infinite. This can also be argued intuitively: in both cases we considered above, the only source of spatial information is the retinal eccentricity of the landmark, but there are infinitely many locations in the environment (along the line connecting the agent with the landmark) from which a landmark is seen exactly at the same retinal eccentricity. (More generally, the two unknown ‘coordinates’ of the agent with respect to the landmark, *d* and *ϵ*, only impact the agent’s observations through the bottleneck of a single variable, retinal eccentricity, and thus one degree of freedom must remain unconstrained.) Mathematically, this can be seen by noting that the uncertainty expressed by the inverse of the Fisher information matrix is scaled by the inverse of the determinant. However, the determinant of the Fisher information matrix is zero in this case:

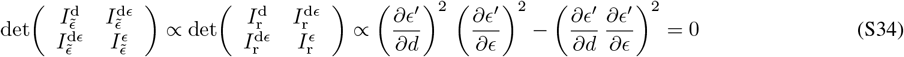

One way to resolve this apparent paradox is to consider an agent that is moving, and the *total* Fisher information accumulated across two consecutive snapshots, separated by a sufficiently small time step. As long as noise (in the observed eccentricity or the retinal responses) is independent across time (at least at the resolution of a time step), the total Fisher information is simply the sum of the Fisher informations that the two snapshots individually provide about the original distance and eccentricity at the time of the first snapshot. Although the total Fisher information matrix is approximately proportional to the Fisher information provided by either snapshot (and therefore the results derived in Eqs. S19, S22, S25, S29, S31 and S33 still apply with a good approximation), its determinant is not zero any more (as long as the retinal eccentricity of the landmark changes):

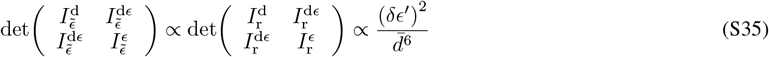

where δ*ϵ*^*′*^ is the change in retinal eccentricity of the landmark between the two snapshots, and 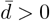 is its geometric mean distance.

Finite uncertainty can also be achieved in a single snapshot, without having to consider motion, if the landmark is not point-like or multiple landmarks are available. Again, the Fisher informations provided by different points on the same landmark (or indeed, by multiple landmarks) approximately add as long as noise (in the observed eccentricity or the retinal responses) is sufficiently independent between them. Therefore, the same mathematical arguments suggest that the determinant of the total Fisher information will be greater than zero, provided that the points (or landmarks) are not co-linear with respect to the agent (i.e. they are seen at different retinal eccentricities).

So far, we have considered *marginal* uncertainties about *d* and *ϵ* – expressing the amount of uncertainty the agent has about either when it only observes retinal eccentricity or responses. For an alternative interpretation that also results in finite uncertainties, one can instead consider *conditional* uncertainty – expressing the amount of uncertainty the agent has about *d* (respectively *ϵ*) when it not only observes retinal eccentricity or responses but also observes, or at least assumes a particular value for, *ϵ* (respectively *d*). In this case, we can make use of the general relationship between a precision matrix (here: the Fisher information matrix), and its associated covariance matrix (i.e. its inverse, here: expressing spatial uncertainty). Given this relationship, the inverse of the individual Fisher information about *d* (respectively *ϵ*) can also be interpreted as the conditional uncertainty (variance) about *d* (respectively *ϵ*). Of course, this conditional uncertainty remains finite (as long as the corresponding individual Fisher information is greater than zero) even in the case of a single snapshot of a single point-like landmark.

### SM4. Bias in distance estimation – analytic intuition

Our numerical simulations of the BION model reproduced different biases in distance estimation between square-vs. trapezoid-shaped environments and between the two halves of a trapezoid environment (Fig. 2d). Here, we provide analytic intuition for the results with a greatly simplified version of the model. In particular, our aim is to explain the seemingly paradoxical effect by which distance estimation biases are *worse* despite homing performance being *better* in the broad than in the narrow side of the trapezoid (Fig. 2d light vs. dark brown bars in the bottom vs. top panels).

For our derivation, we formalize the estimated distance between two target locations 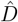 (as measured in the distance estimation task), simply as the distance between two estimated target locations, 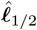(as measured in the homing task). (Note that for the results shown in the main text, Fig. 2, our simulations were performed without making this assumption, following the detailed procedures of the corresponding experiments, see Methods.) Thus, we write

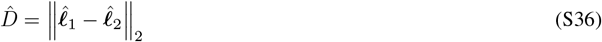

We note that the (square of) the distance trivially decomposes into squared distances along each (x/y) axis

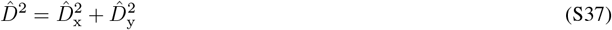

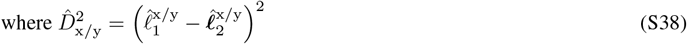

Therefore, in the following, we treat distances and locations along the two axes separately, and drop the x*/*y index to simplify notation. Furthermore, just as when we reasoned about how Eq. 1 predicts the way visual information about location changes in an environment (Fig. 2b, main text), we note that overall information about a target location is dominated by information obtained while looking in the direction of the closest wall at that location. Thus, as a further simplification, for location estimates along each axis, we only consider the case when the agent’s heading direction is aligned with that axis and faces the nearest wall that is orthogonal to that axis.

First, we compute 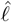 as a function of ℓ, the true location. For this, we take 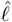 to be the posterior mean with both the prior and visual likelihood assumed to be Gaussians (such that the posterior mean only depends on their means and variances). Let the precision of the prior be

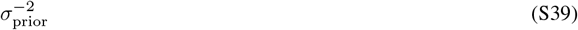

which we expect to depend inversely on the size of the environment along the axis of heading direction. Thus, we expect 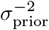 to be smaller along the x-axis than the y-axis of a trapezoid environment, and more specifically, along the y-axis to be smaller in the broad than in the narrow half (Fig. S1a).

Without loss of generality, we take the mean of the prior to be at

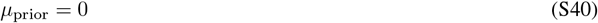

such that the walls at the two ends are respectively at locations −*α L* and (1 − *α*) *L*, with *α* being the relative distance (in units of of the prior mean to the closest wall (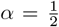 for isotropic environments, such as the square environment, or at least when the heading direction is orthogonal to an axis of symmetry, such as the y-axis in the trapezoid environment; but 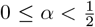 for an axis of asymmetry in an anisotropic environment, such as the x-axis in the trapezoid environment, see also below).

Fig. S1a shows the effects of the anisotropy of the trapezoid environment on the location of the prior mean relative to the walls, the midline of the environment, and the target locations between which distances needed to be estimated in the experiment^1^:

1. The prior mean along the x-axis (Fig. S1a, vertical gray line) is offset towards the broad end, away from broad-narrow boundary (which was defined to be at the midline in the experiment^1^; Fig. S1a, vertical black line), such that it is inside the broad half of the environment (*α* ≈0.39). There is no such offset along the y-axis as the environment is symmetric in that direction (Fig. S1a, horizontal gray line, 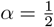).
2. In the experiments^1^, target locations for estimating a given distance were always on the same side of the broad-narrow boundary (Fig. S1a, blue and red dots vs. vertical black line). However, because the prior mean was inside the broad half (Observation 1), target locations in the broad half were often on opposite sides of the prior mean, whereas target locations in the narrow half were always on the same side of the prior mean (Fig. S1a, blue and red dots vs. vertical gray line).
3. There is no shift in the prior along the y-axis (Observation 1), so there is no difference between the broad and narrow sides in whether target locations are on the same or different sides of the prior mean. Specifically, in the experiment^1^, they were typically on opposite sides of the midline (the prior mean; Fig. S1a, blue and red dots vs. horizontal gray line). Given our results about how visual uncertainty scales with environmental geometry (Eqs. 1, S19 and S29, see also Fig. 2a), we take the precision of the visual likelihood to be

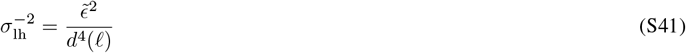

where

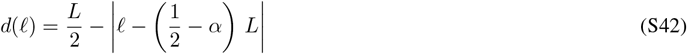

is the distance to the closest wall when the true location of the agent is ℓ ∈ [−*α L*, (1 − *α*) *L*], and

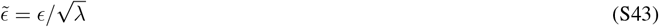

can be treated as the *effective eccentricity* of current visual cues, i.e. the equivalent distance of a single point-landmark from the line of sight that would be as informative as the wall in front of the agent, with *λ* a free parameter controlling the relative informativeness of the prior compared to the likelihood. As such, we expect 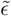 to be greater when the agent moves along the x-axis facing the broad wall than when facing the narrow wall, and even greater when the agent is moving vertically. We approximate the mean of the visual likelihood with the true location

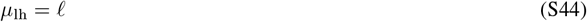

(which is only exact on average, when the precision is sufficiently high). Then, the posterior mean is simply given as the weighted average of the prior and likelihood means, with the corresponding precisions as weighting factors, which can be expressed as:

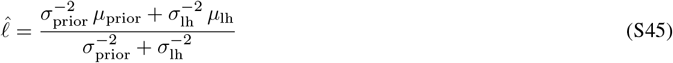

Finally, substituting Eqs. S40, S41 and S44 into Eq. S45, we obtain:

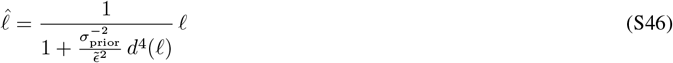

The blue and red solid curves in Fig. S1b-c show 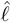 as a function of ℓ along the x- and y-axis of the trapezoid environment in its two halves (broad and narrow). Note that for 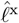 the horizontal extent of the environment defines 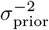 and the vertical extent (that is different in the broad and narrow halves) defines 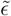 in the equations above (Fig. S1b), and vice versa for 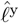 (Fig. S1c). The following properties of the ℓ-to-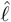mapping are worth noting:
4. In general, 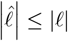, i.e. the curves are all between the 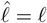 diagonal and the 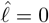 axis. This is because the posterior mean shows regression to the prior mean (at 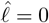 here; Eq. S40).
5. As the precision of the likelihood changes with ℓ (Eq. S41), 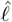 depends nonlinearly on ℓ. (This contrasts with the classical case^66^ in which the precision of the likelihood is constant, and thus 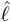 is a linear function of ℓ.)
6. Along the x-axis, the nonlinearity is stronger and overall biases are greater for the narrow than for the broad half (Fig. S1b; the red curve is farther from the diagonal than the blue) – this is because, although the two sides share the same 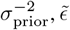 is smaller in the narrow than for the broad half. Importantly, the same is true along the y-axis (Fig. S1c) but for a different reason: now the two halves share the same 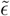, but 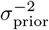 is smaller in the broad than in the narrow half). Note that all of the above refer to (biases in) estimated *locations*. To understand biases in estimated *distances* (our stated goal), it is *differences* in estimated location biases that matter (Eq. S38). Also note that, if visual precision was constant, so that 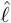 depended linearly on ℓ, a given distance would always be (under)estimated by the same amount, irrespective of where its ‘endpoints’ (the target locations) are. However, because visual precision depends on location and so 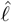 is a nonlinear function of ℓ (Observation 5), the same distance can be estimated with different biases depending on the target locations – and particularly, as we will see below, depending on whether the target locations are in the broad or narrow half of the environment:^*^
7. When the two target locations are on the opposite sides of the prior mean, the distance between them will always be underestimated due to the regression of both estimated locations to the prior mean (Observation 4).
8. When the two target locations are on the same side of the prior mean, due to the nonlinearity of the 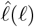 curves (Observation 5), distances can be under-, over-, or even (approximately) accurately estimated depending on the parameters and where each target location lies.
9. Along the x-axis (Fig. S1b), the nonlinearity of location estimates is stronger (Observation 6) and so – depending on the target locations – both the potential under- and overestimation of distances can be more pronounced in the narrow than in the broad half. However, in the broad half, the experimentally used target locations for distance estimation are often on opposite sides of the prior mean, while in the narrow half, they are necessarily on the same side of the prior mean (Observation 2 and Fig. S1a, dots vs. gray vertical line). Therefore, Observation 7 applies (more) in the broad half leading to a consistent underestimation of distances there (Fig. S1d, top, blue), while Observation 8 applies in the narrow half, allowing for a diversity of biases. In particular, with the experimentally used target locations and for a wide range of values for *λ* (the only free parameter in our formulation here), there will be a mixture of over- and under-estimations on the narrow side, and so a less negative (or even positive) average bias on balance (Fig. S1d, top, red).
10. Along the y-axis, location estimates are virtually only biased, if at all, in the broad half (Observation 6; Fig. S1c). In addition, target locations for distance estimates are (almost) always on opposite sides of the prior mean (Observation 3) and so Observation 7 applies. Therefore, distances in the narrow half will be unbiased, and if they are biased at all, they will be underestimated in the broad half (Fig. S1d, middle).
11. In summary, in the broad half, overall distance underestimation dominates along both the x- and y-axes (Observations 9 and 10), and so total distances (Eq. S37) will also be underestimated (Fig. S1d, bottom, blue). In the narrow half, distance estimates are largely unbiased along the y-axis and show a mixture of biases along the x-axis (at least for *λ ≤* 1), and so the latter will also be true for total distances (Fig. S1d, bottom, red). This explains why distance estimation biases are *more* negative in the broad than in the narrow half (Fig. 2d, bottom, light vs. dark brown).

**Fig. S1.**
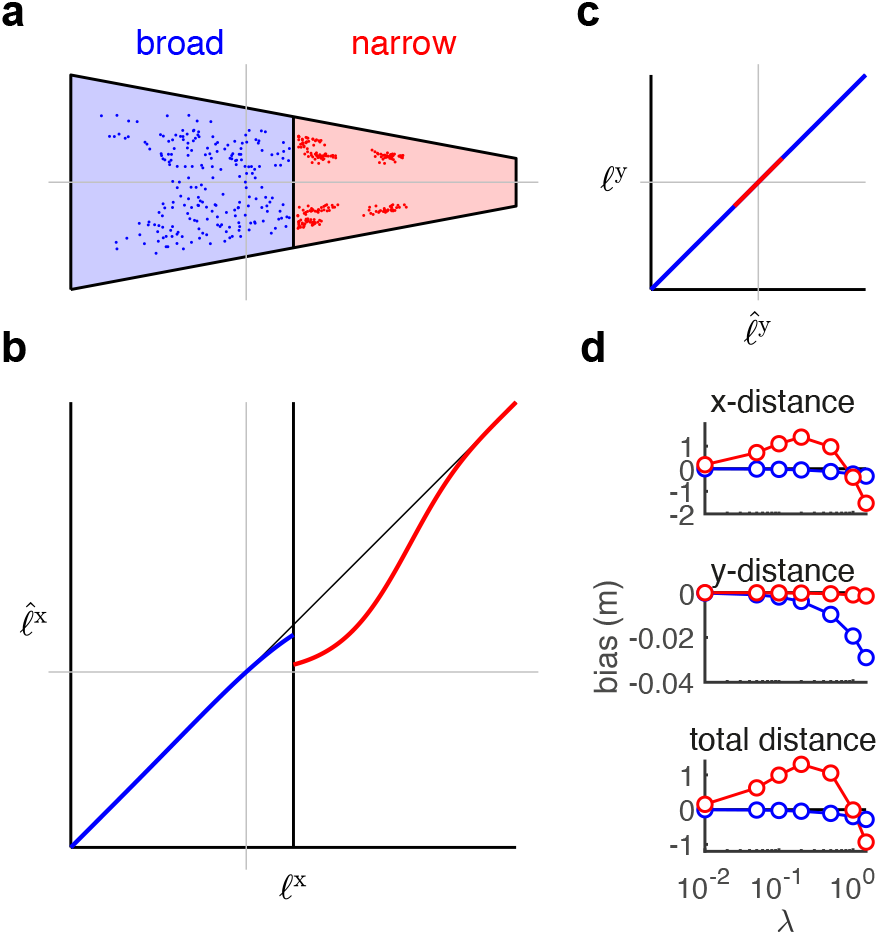
Bias in distance estimation – analytic intuition. **a**. Layout of the trapezoid environment, divided into broad (blue) and narrow halves (red) as defined in the experiment^1^. Gray lines show prior mean along the x- and y-axis, dots show distribution of target locations used for distance estimation in the experiment. **b–c**. Estimated 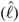 vs. true locations (ℓ) along the x-(**b**) and y-axis of the environment (**c**), in the broad (blue) and narrow half of the environment (red) as given by Eq. S46 with *λ* = 1. (Note that ℓ and 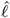 axes are swapped in **c**.) Gray lines show prior mean along the x-(**b**) and y-axis (**c**), on both the ℓ and 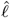 axes of the corresponding panels, vertical black line in **b** shows the broad-narrow boundary of the environment (cf. **a**). **d**. Average biases in distance estimation along the x-axis (top) and y-axis of the environment (middle), and in total (bottom) as a function of *λ*, in the broad (blue) and narrow half of the environment (red). Estimation bias is computed as 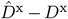 (top), 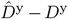 (middle), and 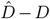 (bottom), where 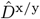 and 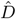are defined as in Eqs. S36-S38, with estimated locations 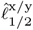 computed as in Eq. S46, *D*^x*/*y^ and *D* are computed analogously from the true locations, 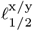, and averages are computed over a distribution of target location-pairs used in the experiment^1^ (cf. dots in **a**). Note that biases are shown with respect to true distances here, whereas Fig. 2d shows biases relative to estimated distances in the square environment (results are virtually identical in the two cases, not shown). To simplify analysis, estimated locations along the x-axis were computed as if the true location along the y-axis was at the midline of the environment (horizontal gray line in **a**), and estimated locations along the y-axis for locations in the broad and narrow half were computed as if the true location along the x-axis was at the broad or narrow end of the environment, respectively (i.e. they represent an upper bound on bias-differences between the two halves).

**Fig. S2.**
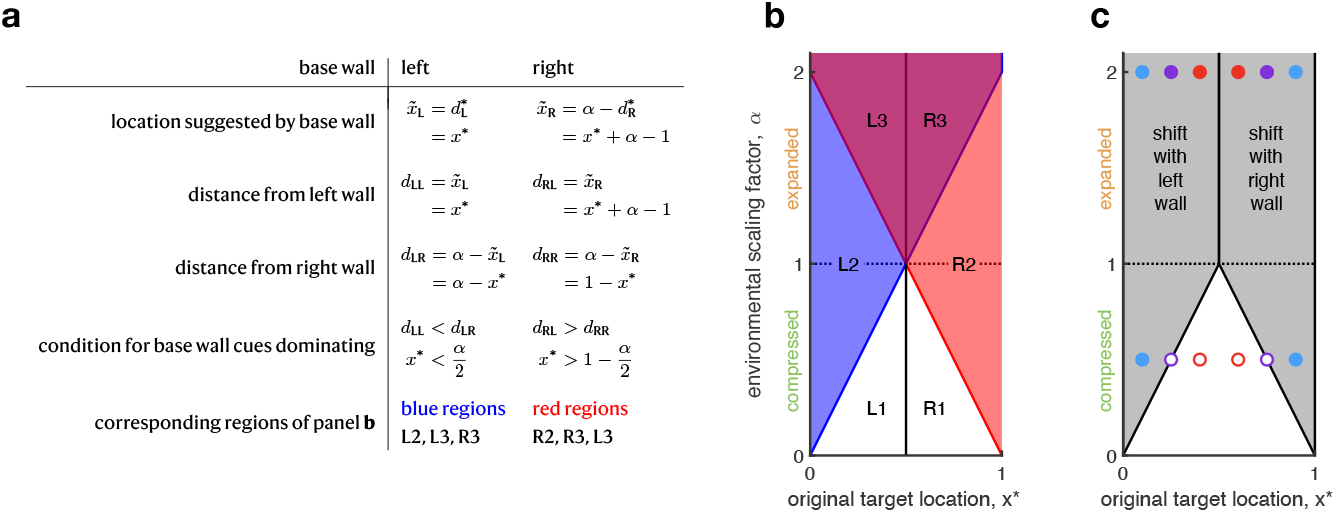
Scaling vs. shifting of homing locations in scaled environments. **a**. Mathematical formulae for scaling vs. shifting of homing locations in scaled environments. 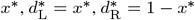, and *α* are the location of the target, its distance from the left and right walls in the original environment (normalized to size 1), and the factor by which the environment is scaled, respectively. See further details in text. **b**. Parameter regions in which the homing locations suggested by neither (L1, R1), either (L2, R2), or both walls (L3, R3) are expressed in behavior. Colored regions are determined by inequalities shown in panel **a**. Purple region is where blue and red regions overlap. **c**. Parameter regions for which shifting of homing locations with the closest wall is predicted in scaled environments (gray areas, based on Eqs. S53 and S54). For comparison, circles show combinations of environmental scaling and target location that were tested experimentally, with filled vs. open circles showing combinations for which shifting vs. scaling was found to dominate. Colors of circles are for illustration only and indicate target locations at small (blue), intermediate (purple), and large distance from the closest wall (red, cf. Fig. 3c-d).

### SM5. Scaling vs. shifting of homing locations in scaled environments – analytic intuition

Our numerical simulations of the BION model correctly predicted that, after environmental scaling, homing responses are best described by shifting with the closest wall when the environment is expanded, and also for targets near the walls when it is compressed, but they are better explained by scaling for targets farther from the walls in compressed environments (Fig. 3c-d). Here, we provide analytic intuition for these results with a greatly simplified version of the model.

In order to obtain a simple intuition, we consider only visual cues in a one-dimensional environment with a point-like landmark at either end. Without loss of generality, we assume that, before scaling, the length of this one-dimensional environment runs from *x* = 0 (the ‘left wall’) to *x* = 1 (the ‘right wall’), and that the right wall is then shifted to *x* = *α* when the environment is scaled (such that 0 *< α <* 1 means the environment has been compressed, and *α >* 1 means it has been expanded). Let the target location before scaling be at 0 *< x*^∗^ *<* 1. Note that at this location, the distances from the left and right walls are 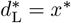 and 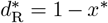, respectively, and so ‘shifting with the closest wall’ happens when homing locations shift with the left wall for *x*^∗^ *<* 1*/*2 and when they shift with the right wall for *x*^∗^ *>* 1*/*2.

In general, during homing, when trying to find one’s way back to the target location, cues based on either the left or the right wall will suggest a homing location that is compatible with the distance of the *original* target location from the corresponding wall. This suggested homing location will thus shift with the corresponding wall if the environment is scaled. However, according to intuitions based on a typical cue combination scenario^67^, homing locations suggested by neither wall would be expected to be expressed purely in behavior because the two sets of cues (those supplied by the left and right wall, respectively) will always be integrated by an ideal observer, such as BION. In turn, the result of this integration would usually be expected to be an interpolation between the two different homing locations – and this interpolation would be detected as ‘scaling’ in the analysis of Fig. 3c-d.

So why does shifting of homing locations occur at all (and in fact in most cases shown in Fig. 3d)? There are two important insights to explain this. First, when Bayesian cue combination occurs between cues of *highly asymmetric informativeness*, although interpolation still happens formally, in practice it will be heavily skewed towards the more informative cue (as the optimal weights for interpolation are proportional to the informative of the cues). Thus, when cues have very asymmetric informativeness, behavior will appear to express only the effect of the dominant (more informative) cue. Second, visual cues in navigation are effectively subject to prominent *signal-dependent noise*: according to Eq. 1, the informativeness of a visual cue for determining the distance from this cue scales inversely with the distance itself. In the following, we show how these two insights explain why shifting rather than scaling of homing responses happens under particular (and in fact most) conditions.

In general, once the environment has been scaled, the location suggested by either wall will be incompatible with cues based on the opposite wall. Under Bayesian cue combination, this location will only be expressed in behavior if the cues from the wall that suggests this location (the base wall) dominate the cues from the opposite wall (first insight above). Given the inverse scaling of the informativeness of cues from either wall with the distance from the corresponding wall (Eq. 1), a suggested location will be expressed in behavior when its distance from the base wall is sufficiently smaller than its distance from the opposite wall (second insight above). Here, we take the simplifying assumption that the shift in dominance between the cues of the two walls is immediate so that cue combination is purely dominated by the closer wall as soon as the observer crosses the midpoint of the (scaled) environment. (We relax this assumption at the end of this section.) Based on these considerations, Fig. S2a shows formulae for the locations suggested by each wall, the distances of these locations from each wall, and the conditions under which these locations are expressed in behavior because the cues of the base wall dominate (illustrated in Fig. S2b).

Note that the results in Fig. S2a-b predict that there will be original target locations for which the homing locations suggested by neither wall (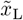or 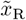 in Fig. S2a) will be purely expressed in behavior – this is where we expect the scaling of homing responses to happen:

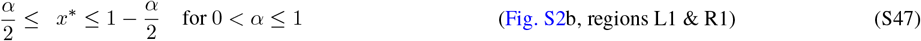

There will also be target locations for which the homing location suggested by only one or the other wall (i.e. either 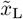 or 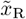) will be eventually expressed in behavior – this will result in pure shifting (Fig. S2b, regions L2 & R2). In particular, 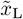 alone will be expressed in behavior when

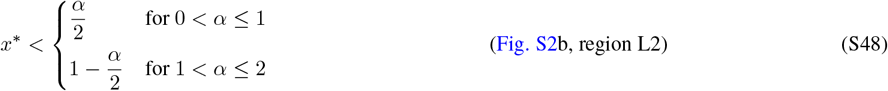

and 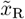 alone will be expressed in behavior when

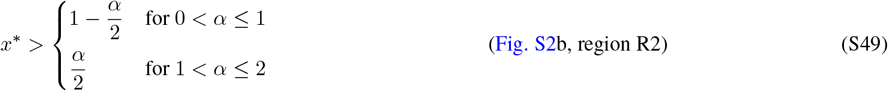

Note that the upper limit on *x*^∗^ in Eq. S48, i.e. for 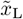 being expressed in behavior, is always smaller than 1*/*2, and the lower limit on *x*^∗^ in Eq. S49, i.e. for 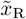 being expressed in behavior, is always greater than 1*/*2. Thus, in each case, the homing location expressed in behavior specifically shifts with the wall to which the original target location was closest.

Finally, there will be a range of target locations for which homing locations suggested by both walls (i.e. both 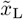 and 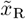) will be expressed in behavior after the environment is expanded:

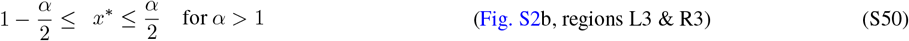

This suggests a bimodal distribution of homing locations, as seen in the full behavior of the model, and also – to some extent – in the data (Fig. 3b). Nevertheless, the experimentally used measure of homing locations in the scaled environment only considers the largest mode of the response distribution^2^ (Fig. 3d). Out of the two modes expressed in behavior the one that is nearer to its base wall will be larger (so 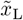 for *x*^∗^ *<* 1*/*2, and 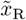for *x*^∗^ *>* 1*/*2). Thus, the base wall with which the largest mode shifts is again the wall that was closest to the original target location, and a shift with the left wall as the closest wall will be measured when

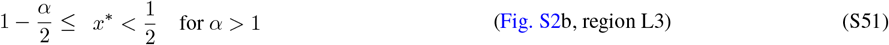

and a shift with the right wall as the closest wall will be measured when

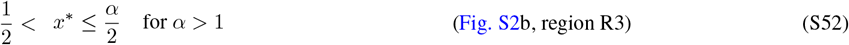

In sum, taking the results in Eqs. S48, S49, S51 and S52 together, homing locations will appear to shift with the closest wall when

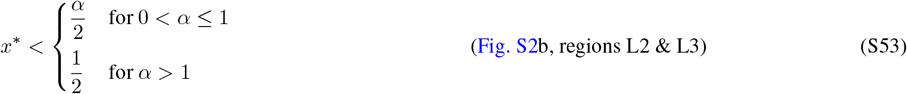

or when

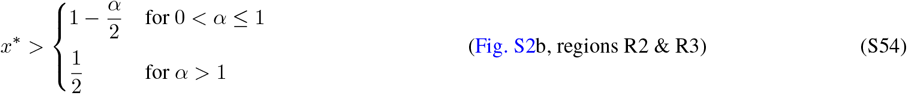

As Fig. S2c shows, Eqs. S53 and S54 mean that homing locations will shift with the closest wall for any *x*^∗^ when the environment is expanded (i.e. when *α >* 1), but they will only shift for *x*^∗^ sufficiently near either wall in compressed environments (*α <* 1) – thus explaining the qualitative behavior seen in humans and exhibited by BION (Fig. 3c-d).

So far, our simple geometry-based analysis assumed that as soon as the distance from one wall becomes even just a little smaller than from the other wall, the informativeness of cues from the two walls become sufficiently strongly asymmetric for cue combination to become totally dominated by the more informative cues of the closer wall and thus to make homing responses shift with that wall. As a slightly better approximation of the full BION model that still allows analytical insight, we can relax this simplifying assumption by considering cue combination with simple Gaussian likelihoods – but at least with a realistic scaling of cue informativeness with distance from the walls (according to Eq. 1). This implies a rapid, but not infinitely fast, shift of dominance towards the closer wall. In line with BION (Eq. 21), we can then compute homing responses as the locations (in the scaled environment) that minimize the ideal observer’s expected mean squared error from the remembered target location (assumed to be at the ground truth target location, *x*^∗^, for simplicity) under its posterior over locations (in the original, unscaled environment) that results from such a combination of cues. The qualitative behavior of such a cue combination model continues to be well predicted by the basic geometric analysis above: scaling of responses happens almost exclusively in compressed environments when the target distance from the closest wall is sufficiently large, and bimodal response distributions arise in expanded environments (not shown).

Conversely, note that the results of our simple geometry-based analysis do not follow trivially from the fact that the shifted equivalents of some original target locations near the middle of the environment (far from the walls) simply fall outside the boundaries of a compressed environment. This simple argument would predict that shifting only becomes impossible once 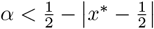 (i.e. the top vertex of the white triangle at the bottom of the plot in Fig. S2c would be at 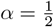 instead of *α* = 1). However, experiments (and the full BION model) show scaling rather than shifting of homing responses for combinations of environmental scaling and target location that do not satisfy this condition (Fig. S2c, open circles are at 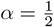 but 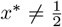).

## Footnotes

* When the two target locations between which distance needs to be estimated are (near-)identical along a given axis, i.e their true distance is zero along that axis, the biases in their estimates are going to cancel, and so their distance estimate will be unbiased (zero) and, as such, it will not contribute to average biases in distance estimates as measured experimentally. Thus, here we ignore these cases.

